# Genome Sequencing and Transcriptome Analysis Reveal Recent Species-specific Gene Duplications in the Plastic Gilthead Sea Bream

**DOI:** 10.1101/769240

**Authors:** Jaume Pérez-Sánchez, Fernando Naya-Català, Beatriz Soriano, M. Carla Piazzon, Ahmed Hafez, Toni Gabaldón, Carlos Llorens, Ariadna Sitjà-Bobadilla, Josep A. Calduch-Giner

## Abstract

Gilthead sea bream is an economically important fish species that is remarkably well-adapted to farming and changing environments. Understanding the genomic basis of this plasticity will serve to orientate domestication and selective breeding towards more robust and efficient fish. To address this goal, a draft genome assembly was reconstructed combining short- and long-read high-throughput sequencing with genetic linkage maps. The assembled unmasked genome spans 1.24 Gb of an expected 1.59 Gb genome size with 932 scaffolds (∼732 Mb) anchored to 24 chromosomes that are available as a karyotype browser at www.nutrigroup-iats.org/seabreambrowser. Homology-based functional annotation, supported by RNA-seq transcripts, identified 55,423 actively transcribed genes corresponding to 21,275 unique descriptions with more than 55% of duplicated genes. The mobilome accounts for the 75% of the full genome size and it is mostly constituted by introns (599 Mb), whereas the rest is represented by low complexity repeats, RNA retrotransposons, DNA transposons and non-coding RNAs. This mobilome also contains a large number of chimeric/composite genes (i. e. loci presenting fragments or exons mostly surrounded by LINEs and *Tc1/mariner* DNA transposons), whose analysis revealed an enrichment in immune-related functions and processes. Analysis of synteny and gene phylogenies uncovered a high rate of species-specific duplications, resulting from recent independent duplications rather than from genome polyploidization (2.024 duplications per gene; 0.385 excluding gene expansions). These species-specific duplications were enriched in gene families functionally related to genome transposition, immune response and sensory responses. Additionally, transcriptional analysis of liver, skeletal muscle, intestine, gills and spleen supported a high number of functionally specialized paralogs under tissue-exclusive regulation. Altogether, these findings suggest a role of recent large-scale gene duplications coupled to tissue expression diversification in the evolution of gilthead sea bream genome during its successful adaptation to a changing and pathogen-rich environment. This issue also underscores a role of evolutionary routes for rapid increase of the gene repertoire in teleost fish that are independent of polyploidization. Since gilthead sea bream has a well-recognized plasticity, the current study will advance our understanding of fish biology and how organisms of this taxon interact with the environment.

## Introduction

Gilthead sea bream (*Sparus aurata*) is a temperate marine coastal finfish that belongs to the Sparidae family, order Perciformes. It is an economically important species highly cultured throughout the Mediterranean area with a yearly production of more than 218,000 metric tonnes, mostly concentrated in Turkey, Greece, Egypt and Spain (FAO, FishStat database, 2019). This species occurs naturally in the Mediterranean and the Eastern Atlantic Seas, from the British Isles and Strait of Gibraltar to Cape Verde and Canary Islands, supporting previous studies of genetic structure a strong genetic subdivision between Atlantic and Mediterranean populations (Alarcón et al., 2004; De Inocentiis et al., 2004). Intriguingly, strong subdivisions have also been found at short distances along the Tunisian coasts (Ben Slimen et al., 2004) or between the French and Algerian coasts (Chaoui et al., 2009). However, unconstrained gene flow occurs along the coast of Italy, in the absence of physical and ecological barriers between the Adriatic and Mediterranean Seas (Franchini et al., 2012).

Gilthead sea bream is a protandrous hermaphrodite species, as it matures as male during its first and second years, but most individuals change to females between their second to fourth year of life (Zohar et al., 1978). This sexual dimorphism is a fascinating subject in evolutionary biology, and Pauletto and coworkers (2018) showed for the first time in a hermaphrodite vertebrate species that the evolutionary pattern of sex-biased genes is highly divergent when compared to what is observed in gonochoristic species. Adaptation to varying environments, including high tolerance to changes in water salinity, dissolved oxygen concentration, temperature, social hierarchy or diet composition are also a characteristic feature of gilthead sea bream, making this species a rather unique fish with a high plasticity to farming and challenging environments. This has been assessed in a number of physiological studies with focus on nutrition (Benedito-Palos et al., 2016; Simó-Mirabet et al., 2018; Gil-Solsona et al., 2019), chronobiology (Mata-Sotres et al., 2015; Yúfera et al., 2017), feeding behavior (López-Olmeda et al., 2009; Sánchez et al., 2009), stress (Calduch-Giner et al., 2010; Castanheira et al., 2013; Pérez-Sánchez et al., 2013; Bermejo-Nogales et al., 2014; Magnoni et al., 2017; Martos-Sitcha et al., 2017; Martos-Sitcha et al., 2019) or disease resilience (Cordero et al., 2016; Estensoro et al., 2016; Piazzon et al., 2018; Simó-Mirabet et al., 2018). However, the underlying genetic bases of this adaptive plasticity remain unknown.

In addition to the two rounds of whole genome duplication (WGD) that affected bony vertebrates (Dehal and Boore, 2005), a third event of WGD (3R) occurred in the genome of the ancestor of teleost fish that is still present in the signature of modern teleost genomes (Jaillon et al., 2004; Kasahara et al., 2007). More recent WGD events occurred at the common ancestor of cyprinids and salmonids (Macqueen et al., 2014; Chen et al., 2019). Comparative genomic analyses have shown that, generally, WGDs are followed by massive and rapid genomic reorganizations driving the retention of a small proportion of duplicated genes (Langham et al., 2004). However, recent studies in rainbow trout (*Oncorhynchus mykiss*) reveal that the rediploidization process can be stepwise and slower than expected (Berthelot et al., 2014). Further complexity comes from tandemly-arrayed genes that are critical zones of adaptive plasticity, forming the building blocks for more versatile immune, reproductive and sensory responses in plants and animals including fish (Rizzon et al., 2006; Kliebesntein 2008; van der Aa et al., 2009; Lu et al., 2012). In any case, it has been shown that retained genes following WGDs or small scale duplicates are preferentially associated with species-specific adaptive traits (Maere et al., 2005). This notion is reinforced by the recently published study of large-scale ruminant genome comparisons (Chen et al., 2019), also evidenced in the case of modern teleosts and primitive eels (Chen et al., 2008; Tine et al., 2014; Rozenfeld et al., 2019) for their improved adjustment to natural environment.

Here we produced a high quality draft sequence of the gilthead sea bream genome by combining high-throughput sequencing with genetic linkage maps. The current draft assembly spans ∼1.24 Gb with 932 scaffolds ordered and oriented along 24 chromosomes derived from the genetic linkage map of the first gilthead sea bream genome release (Pauletto et al., 2018). Homology-based functional annotation, supported by RNA-seq transcripts, identified 55,423 actively transcribed genes corresponding to 21,275 unique descriptions. Synteny and phylogenomic analyses revealed a high frequency of species-specific duplications, mostly resulting in the enrichment of biological processes related to genome transposition but also to immune response and sensory responses. Since divergent regulation and function of the multiple copies of tissue-exclusive genes is also supported by RNA-seq transcriptional analysis, gilthead sea bream is emerging as an interesting model to assess the teleost genome expansion and its contribution to adaptive plasticity in a challenging environment.

## Material and methods

### Ethics Approval

Procedures for fish manipulation and tissue collection were approved by the Ethics and Animal Welfare Committee of Institute of Aquaculture Torre de la Sal and carried out according to the National (Royal Decree RD53/2013) and the current EU legislation (2010/63/EU) on the handling of experimental fish.

### Fish and Tissue Processing

Fish were reared from early life stages under natural conditions of photoperiod and temperature at the experimental facilities of IATS (40°5N; 0°10E). Blood of one single male was obtained from caudal vessels using heparinized syringes, and DNA from total blood cells was extracted with a commercial kit (RealPure Spin Blood Kit, Durviz, Valencia, Spain). Quality and quantity of genomic DNA was assessed by means of PicoGreen quantification and gel electrophoresis. An aliquot of 5 µg DNA was mechanically sheared with a bath sonicator (Diagenode BioRuptor, Diagenode, Liège, Belgium) and low molecular weight fragments were used for the preparation of DNA libraries.

Total RNA (70-100 µg) from white skeletal muscle (6 individual fish) and pooled samples of anterior and posterior intestine sections were extracted with the MagMAX™-96 Total RNA Isolation Kit (Applied Biosystems, Foster City, CA, USA). The RNA concentration and purity was determined using a Nanodrop 2000c (Thermo Scientific, Wilmington, DE, USA). Quality and integrity of the isolated RNA were checked on an Agilent Bioanalyzer 2100 total RNA Nano series II chip (Agilent, Amstelveen, Netherlands), yielding RNA integrity numbers (RIN) between 8 and 10.

### DNA/RNA Sequencing

Genomic DNA material was used for the preparation of two standard TrueSeq Illumina libraries (Illumina Inc) with an average size of 360 and 747 bp, respectively. Illumina NextSeq500 system under a 2×150 paired-end (PE) format was used as sequencing platform to generate approximately 600 million reads. Additionally, two different strategies were implemented in order to help in genome scaffolding: 1) Nextera Mate-Pair Preparation Kit (Illumina Inc) was used to make two mate pairs (MP) libraries (average insert sizes were 5 and 8 kb) using the Illumina NextSeq500 platform to a depth of 11 Gb (2×75 MP format) and 2) genomic DNA was submitted to Macrogen (Seoul, South Korea) for the construction of 12 single molecule real time (SMRT) cell libraries (insert size up to 50 kb) using PacBio RS II (Pacific Biosciences) as sequencing system. Additionally, eight RNA-seq libraries (for more details, see Data Availability) were constructed by means of Illumina TrueSeq RNA-seq preparation protocol (non-directional method). Sequencing of indexed libraries was performed on the Illumina Hiseq v3, resulting in approximately 11-17 million reads per sample (1×75 nt single reads) from skeletal muscle samples and 22-27 million read pairs (2×150 nt paired reads) from intestine samples.

### *De novo* Genome Assembly and Chromosome Anchoring

The SMRT cell libraries were pre-processed using the trimming of the CANU assembler (Koren et al., 2017). Illumina PE libraries were checked for quality analysis using FASTQC 0.11.7, available at (http://www.bioinformatics.bbsrc.ac.uk/projects/fastqc/), and then pre-processed using Cutadapt v1.16 (Martin, 2011) and Prinseq 0.20.4 (Schmieder and Edwards, 2011). Quality analysis and pre-processing of Illumina MP libraries was performed with FastQC and Platanus (Kajitani et al., 2014). These protocols for pre-processing *de novo* assembly were executed using the DeNovoSeq pipeline provided by the GPRO suite (Futami et al., 2011). Jellyfish (Marçais and Kingsford, 2011) was used to estimate the genome size calculating the count distribution of k-mers in the set of Illumina PE libraries. The estimated coverage was inferred using Bowtie2 v2.3.4.1 (Langmead et al., 2009). Illumina PE and MP libraries were introduced in the 127mer version of the assembler SOAP de Novo2 v2.04-r241 (Luo et al., 2012) for the assembly of gilthead sea bream genome. In order to test different k-mer values, different assemblies were performed and a k-mer length of 63 bp (k63) was considered the best in terms of metrics. To improve the consensus sequence and to close gaps, two rounds of the following combined strategy were conducted: 1) elimination of duplicates with Dedupe of BBTools (http://jgi.doe.gov/data-and-tools/bbtools/), 2) gap filling using PacBio corrected reads with PBJelly (English et al., 2012), 3) gap filling using PE and MP libraries with Soap *de novo* Gap Closer, 4) hybrid re-scaffolding using corrected SMRT reads together with Illumina PE and MP reads with Opera 2.0.6 (Gao et al., 2011) and 5) transcriptome-guided re-scaffolding using as reference the gilthead sea bream transcriptome (Calduch-Giner et al., 2013) with L RNA scaffolder (Xue et al., 2013). A step of genome masking was not considered in order to achieve a more reliable genome draft.

Highly conserved non-coding elements (CNEs) present in 3 hermaphrodite genomes (*S. aurata, Lates calcarifer, Monopterus albus*) were released by Pauletto et al., (2018), and the super-scaffold coordinates related to these CNEs (200-800 bp interval length) were then retrieved. Sequences were aligned against our assembly for increasing the super-scaffolding by means of the BLAST package. A genome browser was built for the navigation and blast-query of the assembled sequences and associated annotations using Javascript-based tool JBrowse (Skinner et al., 2009). The genome browser, available online at http://nutrigroup-iats.org/seabreambrowser, provides two modes of navigation for the assembly scaffolds and the entire set of super-scaffolds anchored from CNEs.

### Genome Annotation

Prediction of coding genes was carried out using the software AUGUSTUS 3.3 in a two-step process. An initial round of prediction was conducted, and gene model parameters were trained from a set of 13 fish species (*Astyanax mexicanus*, *Danio rerio*, *Gadus morhua*, *Gasterosteus aculeatus*, *Latimeria chalumnae*, *Lepisosteus oculatus*, *Oreochromis niloticus*, *Oryzias latipes*, *Petromyzon marinus*, *Poecilia formosa*, *Takifugu rubripes*, *Tetraodon nigroviridis* and *Xiphophorus maculatus*) available in the Ensembl database release 87 (Cunningham et al., 2015). Then, the merged prediction of gilthead sea bream genes was translated to peptides using OrfPredictor script (Min et al., 2013), and it was used by Scipio 1.4 (Keller et al., 2008) to generate a new training set for a second round of gene predictions. This second round included sequences from the published gilthead sea bream transcriptome (Calduch-Giner et al., 2013) and RNA-seq data from muscle and intestine in addition to those of liver, gills and spleen, retrieved from the SRA archive (see Data Availability) (Piazzon et al., 2019) as AUGUSTUS hints. The script autoAugTrain.pl of AUGUSTUS was used to determine the precise exon/intron gene structures. The Gffread software (Trapnell et al., 2012) rendered the final set of coding sequences (CDS), using the genome transcript file generated by AUGUSTUS. BLAST package was used for gene annotation, performing BLASTX searches against SWISSPROT, NR and the IATS-CSIC gilthead sea bream transcriptome databases with an E-value cutoff of 10^-5^ using the DeNovoSeq pipeline provided by the GPRO suite. Redundancy analysis were performed in order to detect segmental duplications (i.e. predicted genes that occur at more than one site within the genome and typically share >90% of sequence identity) within the final set of transcripts retrieved from RNA-seq libraries using Dedupe of BBTools (http://jgi.doe.gov/data-and-tools/bbtools/). Identity thresholds in redundancy analysis were fixed at 90%, 95% and 98%.

The mobilome draft was annotated considering the following mobile genetic elements (MGEs): non-coding RNA genes, introns, low complexity repeats, Class I retrotransposons, Class II DNA transposons and Chimeric/Composite genes. Introns were retrieved from the *ab initio* predictions. To annotate non-coding RNAs (ncRNAs), a non-redundant database of both small and long ncRNAs was constructed based on the ncRNAs annotations of fish genomes used for *de novo* gene prediction (Maere et al., 2005). An additional fish tRNA database was created using the tRNAs from *D. rerio*, *G. aculeatus*, *O. latipes*, *P. marinus*, *T. rubripes* and *T. nigroviridis* from UCSC (http://gtrnadb2009.ucsc.edu). Then, a BLAT search (Kent, 2002) served to annotate ncRNAs in the gilthead sea bream genome. Duplicated BLAT outputs were removed using Bedtools (http://bedtools.readthedocs.io). A final step of curation was performed based on the merging of entries that in the same scaffold had: 1) the same parent and were consecutive in 5-10 nucleotides, 2) the same target and initial position), 3) the same biotype and overlapped and 4) the support of real transcripts from the gilthead sea bream transcriptome. After curation, repeat sequences retained into longer ones were discarded. To annotate the remaining MGEs, RepeatModeler 1.0.11 (www.repeatmasker.org) was used for the *de novo* repeat family identification. RepeatMasker 4.0.7 and NCBI-BLAST alignments (E-value threshold < 10^-5^) (Altschul et al., 1990) were used to identify simple repeats, low complexity repeats and interspersed repeats within the gilthead sea bream genome. Repbase 22.09 (Bao et al., 2015), GyDB (Llorens et al., 2011) and *de novo* repeat families coming from RepeatModeler were used as libraries. LTR finder (Xu and Wang, 2007) and Einverted of EMBOSS (Rice et al., 2000) were used to characterize long terminal repeats (LTRs) and inverted repeats, respectively.

All the annotations corresponding to coding genes associated to MGEs (chimeric/composite genes) were extracted from the previously presented annotation of coding gene and were used as queries in a BLAST search against Repbase 22.09 and GyDB databases. All the results were curated by means of merging overlapping features with the same annotation or separated by less than 100 nucleotides.

### Gene Synteny and Phylogenomics

Synteny detection was performed across the genome of gilthead sea bream over other 9 fish species (*Cynoglossus semilaevis, D. rerio, G. aculeatus, Maylandia zebra, O. mykiss, O. niloticus, O. latipes, Salmo salar and Xiphophorus maculatus)*. The algorithm includes the following steps: 1) selection of single-copy genes present in only one scaffold in the gilthead sea bream assembly, 2) alignment of gilthead sea bream genes against the other species with BLASTX of the NCBI-BLAST package with more than 70% of sequence identity and coverage, and 3) synteny file construction, establishing an E-value < 10^-5^ to consider a gilthead sea bream-species gene correspondence (with number of gaps < 25). A syntenic block must contain a minimum of 5 genes to be included in the results. Circular genome representations were created using Circos (Krzywinski et al., 2009).

The gilthead sea bream phylome was reconstructed using phylomeDB pipeline (Huerta-Cepas et al., 2014). For each protein-coding gene in gilthead sea bream, a Smith-Waterman search was performed against the proteome database of 19 selected species (*Latimeria chalumnae, L. oculatus, D. rerio, A. mexicanus, P. formosa, G. morhua, O. mykiss, Scophthalamus maximus, O. latipes, O. niloticus, T. rubripes, G. aculeatus, T. nigroviridis, Petromyzon marinus, Callorhinchus milii, Xenopus traevis, Mus musculus and Anolis carolinensis*). Multiple alignments of homologous sequences (E-value < 10^-5^ and 50% overlap over query sequence) were built in forward and reverse sense with three sequence alignment programs: MUSCLE (Edgar, 2004), MAFFT (Katoh et al., 2005) and KALIGN (Lassman and Sonnhammer, 2005). The six resulting alignments were then combined in a consistency framework as implemented in M-COFFEE (Wallace et al., 2006), and the resulting alignment was trimmed with trimAl (consistency cut-off of 0.16667 and -gt > 0.1) (Cappella-Gutiérrez et al., 2009). Multiple trees were then built, and the programming toolkit ETE (Huerta-Cepas et al., 2010) was used for each tree to understand duplication and speciation relationships by means of a 0-score species overlap approach. All information about orthology and paralogy relationships is available in phylomeDB (Huerta-Cepas et al., 2014). Gene duplication in the gilthead sea bream lineage was analyzed to detect genes that had undergone duplications through the evolution in different lineages (Huerta-Cepas and Gabaldon, 2011). PhyML v3 (Guindon et al., 2010) was used to create a maximum likelihood tree with one-to-one orthologous in each of the selected species. Branch support was analyzed using a parametric approximate likelihood ratio test (aLRT) based on a chi-square distribution with three rates categories in all the cases. A super-tree from all single gene trees in the gilthead sea bream phylome was also reconstructed using a gene tree parsimony strategy as implemented in duptree (Wehe et al., 2008).

### Functional Gene Enrichment Analysis

A functional analysis of gene ontology (GO) terms and metabolic pathways was performed over the protein coding genes (PCG) model. Cellular Component, Molecular Function and Biological Process GO terms were obtained from this functional analysis and a threshold of 50 counts was used to achieve the most representative GO terms for each category. Fisher test-based functional enrichment of biological process-associated GO terms was computed by analysing the fraction of the model corresponding to chimeric/composite genes. Enrichment analysis derived from phylogenomics was also performed using FatiGO (Al-Sharour et al., 2007) by comparing ontology annotations of the proteins involved in duplication against all the others encoded in the genome.

### Gene Duplication Landscape and Tissue Gene Expression

RNA-seq sequenced reads were processed to generate a gene expression Atlas across tissues. Briefly, reads were independently mapped against the reference transcriptome created from the set of *ab initio* predictions using Bowtie2. As a highly conservative procedure, only predictions with > 50% homology overlapping and ≥ 5 counts were accepted and included as reliable features. Corset v1.07 (Davidson and Oshlack, 2014) was used to quantify genes in each sample separately. Expression values were calculated in reads per kilobase per million mapped reads (RPKM) (Mortazavi et al., 2008).

To retrieve and annotate duplication events, we considered both the species-specific set of homologous genes from the phylogenomics analysis as well as *ab initio* predictions supported by RNA-seq transcripts. To consider a tissue-specific set of paralogs, all the copies must be supported by phylogenomic evidence and showing the same molecular description based on sequence similarity. Furthermore, to consider a tissue-exclusive set of paralogs, all the copies must also show an expression value in only one of the analyzed tissues. A statistical t-test and a one-way ANOVA (P < 0.05) test were used to detect the differential expression between specialized gilthead sea bream paralogs of skeletal muscle, liver, gills and spleen. Correction by False Discovery Rate (FDR) (α = 0.05) was applied for all the paralog sets. This statistical analysis was not applied to intestine samples because the expression analysis was conducted with pooled instead of individual samples.

The existence of Atlas of expression in humans and other higher vertebrates (https://www.proteinatlas.org, https://www.ebi.ac.uk/gxa/home) was exploited to retrieve and compare the enrichment of tissue-exclusive paralogs. Accordingly, tissue-exclusive genes with non-redundant descriptions (initially assessed by RNA-seq) were categorized as follows: 1) enriched genes in the same tissue in other animal models, 2) enriched genes in the same tissue and in other tissues present in the analysis, 3) genes expressed in almost all the analyzed tissues and 4) unclassified genes.

### Real-time qPCR Validation

Duplicated genes from the analyzed tissues, covering a wide range of expression level among copies, were chosen for real-time qPCR validation: *cav3*, *myod1* and *myod2* (skeletal muscle); *slc6a19* and *aoc1* (intestine); *upp2* and *prom1* (liver); *lmo1* and *yjefn3* (gills); *gp2* and *hbb2* (spleen). Genbank accession numbers of the aforesaid duplicated transcripts are MN131091-MN131112. To complete the range of expression, *cdh15* (skeletal muscle), *cldn15* (intestine), *clec10a* (liver), *sox3* (gills) and *lgals1* (spleen) were included in the qPCR. The validation was performed on the same RNA individual samples used for RNA-seq. Primer design (Supplementary Table 1), reverse transcription, qPCR optimization and reactions were performed as previously detailed (Benedito-Palos et al., 2016). Specificity of reactions was verified by melting curves analyses and expression data were normalized to *β-actin* using the delta delta Ct method (Livak and Schmittgen, 2001). Pearson correlation coefficients were calculated in order to compare gene expression values for RNA-seq samples and qPCR expression data.

## Results

### Reads Sequencing Reveals a Large Genome Size

Gilthead sea bream genome was assembled using a hybrid strategy involving Illumina NextSeq500 and PacBio RS II as sequencing platforms. An overview of the main stages and achievements of the project is shown in Figure 1. Data obtained from the two PE and two MP Illumina libraries reached ∼94.8 Gb and ∼11.7 Gb, respectively (see Supplementary Table 2). PE read assembly yielded 51,918 contigs with an N50 of 50.2 kb and an L50 of 6,823 contigs. The initial assembly was further improved by means of scaffolding with MP and SMRT reads followed by gap filling. This procedure resulted in 5,039 scaffolds (>750 bp length) with an N50 scaffold length of 1.07 Mb and an L50 scaffold count of 227. At this end, the percentage of assembly in scaffolded contigs was 99.2% with a mean scaffold size of 247.38 kb and an average GC content of 39.82%. For more details in assembly metrics see Supplementary Table 3.

**Figure 1.**
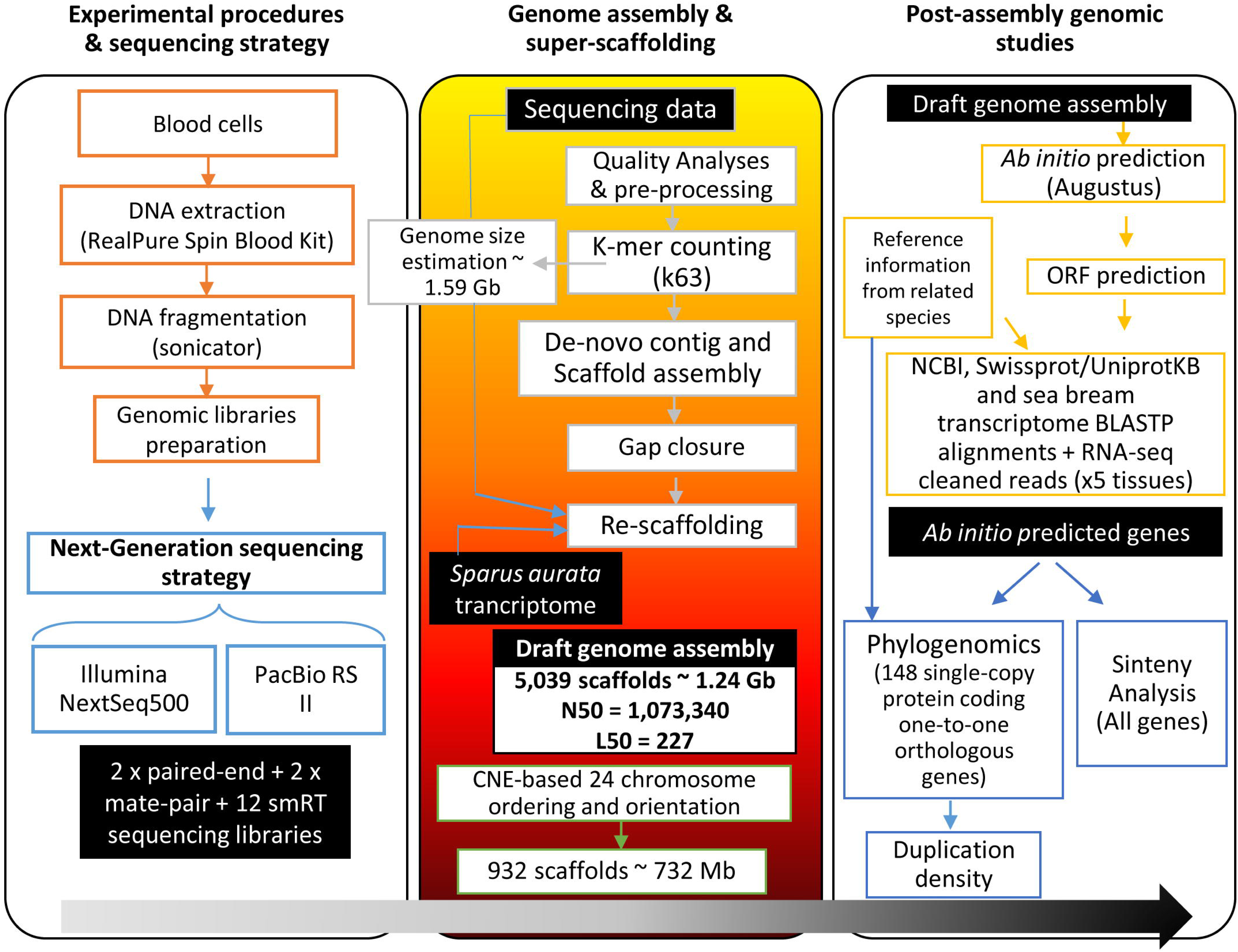
Workflow of the gilthead sea bream genome assembly project. Black boxes with white text indicate generated genomic resources, according to the following steps: experimental procedures & sequencing, genome assembly & super-scaffolding, and post-assembly analyses over the genome draft (*ab initio* gene prediction, synteny analysis, phylogenomics).

K-mer analysis using PE reads (Supplementary Figure 1A) showed 63-mer read length frequency with an estimated genome size of ∼1.59 Gb (main peak), including 543 Mb of repeated k-mers (repeat peak). The total scaffold length was ∼1.24 Gb, which represents 78% of the estimated total genome size. According to this, the average assembly coverage was 67.8x, and 90% of the total assembled genome was included in the largest 1,613 scaffolds (Supplementary Figure 1B).

Super-scaffolding assembly was performed using 7,700 CNEs derived from the genetic linkage map of the first gilthead sea bream genome release (Pauletto et al., 2018). These CNEs, associated to unique positions within 932 scaffolds, served for ordering and orienting 57.8% of the scaffold assembly length (∼732 Mb) in 24 super-scaffolds (Supplementary Figure 2). The resulting virtual gilthead sea bream karyotype can be viewed at www.nutrigroup-iats.org/seabreambrowser.

### Multiple Gene Duplications Are Surrounded by Transposable Elements

A first *ab initio* prediction of PCG was carried out using AUGUSTUS v3.3 (Stanke et al., 2008). To support the establishment of the PCG model, eight RNA-seq libraries from this study (6 skeletal muscle, 2 intestine) in combination with additional libraries from liver (4), spleen (3) and gills (3) (retrieved from SRA archive) were processed to generate an Atlas of gene expression across tissues (see accession numbers in Data Availability). The sequenced reads were mapped against *ab initio* predictions, and 55,423 PCG were inferred based on RNA-seq transcriptome analysis and homology against SWISSPROT, NR or the IATS-CSIC gilthead sea bream transcriptome database (Calduch-Giner et al., 2013). This procedure generated a total of 21,275 unique gene descriptions with 9,250 single-copy genes. Up to 90% of unique gene descriptions are comprised in the 1,613 largest scaffolds (Figure 2A). The average gene length is 10,134 bp with exon and intron mean sizes of 184 bp and 1,751 bp, respectively. This yields an average protein length of 375 amino acids. For super-scaffolded genes, the number of non-redundant protein descriptions decreases to 16,046 with an average gene size of 11,756 bp (Figure 2B). Dedupe redundancy analysis performed over the transcript set retrieved from RNA-seq revealed a total of 559 duplicated genes, which represents a small fraction (1.01%) of segmental gene duplications (Supplementary Table 4). Furthermore, the number of containments (i.e. shorter overlapping contained sequences) at 98%, 95% and 90% of identity threshold was also very low (3.31%, 5.05% and 6.83%, respectively).

**Figure 2.**
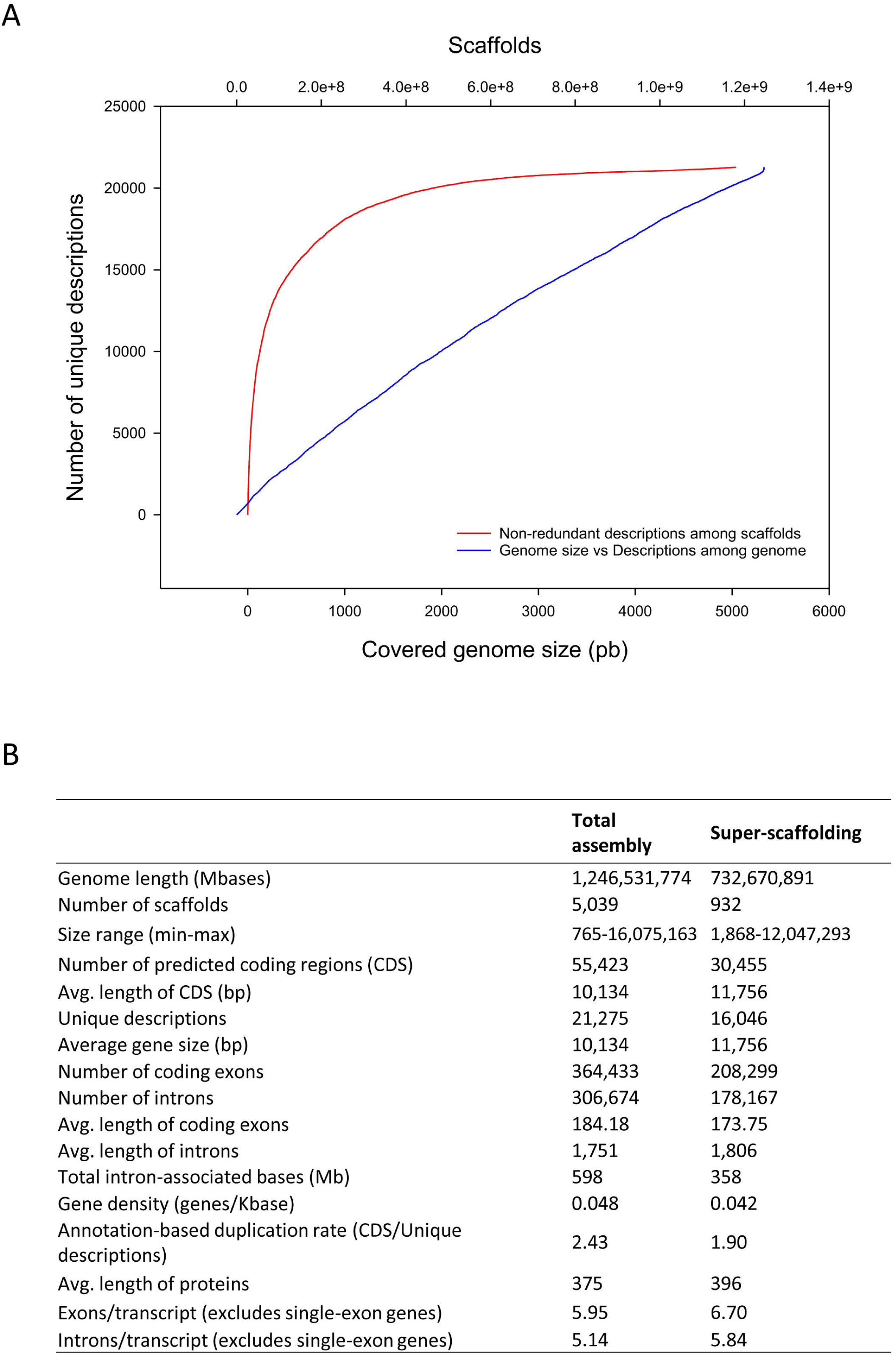
Scaffold unique descriptions distribution and gene features. **(A)** Cumulative distribution of non-redundant gene annotations among length-ordered scaffolds. **(B)** Summary statistics of gene annotation in the gilthead sea bream genome.

At the scaffold level, the gilthead sea bream mobilome accounts for the 75% of the full genome size (944 Mb). More than 60% of this mobilome (599 Mb) is constituted by introns, whereas the rest of MGEs are widely spanned throughout the assembly (Supplementary Table 5). The predicted low complexity repeats (16.91%) spanned 160.5 Mb with approximately 160 Mb corresponding to 2,500 repeat families classified as *de novo* specific of gilthead sea bream. The remaining 0.5 Mb corresponded to known repeats (inverted and/or tandem repeats as well as satellites and microsatellites) also present in other fish genomes. Class I MGE (5.84%) comprised 27.2 Mb of LTRs retroelements (*Ty3/Gypsy, BEL/Pao, Ty1/Copia* and *Retroviridae*-like), 27.8 Mb of non-LTR retroelements (distributed in 14 families, mainly LINEs and SINEs), and 0.2 Mb of YR-like DIRS retrotransposons. Class II MGE (10.55%) included 99.6 Mb split in 27 groups of DNA transposons (mainly *hAT, Tc1/mariner, PIF/Harbinger* and *PiggyBac* elements). The last fraction of the mobilome corresponded to non-coding RNA (1.25%) and chimeric/composite genes (1.95%). A complete list of non-coding RNA (ncRNA) genes is shown in Supplementary Table 6, including both long (11 Mb constituted by 10 groups; mainly lincRNA, pseudogenes and processed transcripts) and small (1 Mb split in 11 groups mainly microRNA, tRNA and snoRNA) ncRNA. Chimeric/composite genes (i.e. those carrying exon traits constituted by MGEs) were split in 10 groups of loci: non-LTR retroelement traits (7 Mb), LTR retroelement traits (0.7 Mb), DNA transposon traits (5.8 Mb), ncRNA gene traits (0.053 Mb), repeats (0.001 Mb), viral-related traits (0.2 Mb) and YR retroelement traits (0.02 Mb), as well as clan AA peptidases (0.047 Mb), Scan/Krab genes (0.008 Mb) and unknown genes (4 Mb). For more specific details about chimeric/composite gene annotation see Supplementary Table 7. Krona representation of split sublevels of mobilome can be seen in Supplementary Figure 3.

### Chimeric Genes Enriched in Immune Response and Response to Stimulus Processes

Functional annotation of gilthead sea bream genes using GO resulted in a diverse set of functional categories allocated to 43,221 genes (Cellular Component, 41,423; Molecular Function, 38,505; Biological Process, 38,588). The top 12 categories of each ontology for non-redundant protein descriptions are shown in Fig. 3A. Cellular component GO terms had the higher gene count with cytoplasm (GO:0005737; 20,689), plasma membrane (GO:0005886; 16,138) and integral to membrane (GO:0016021; 12,436) GO terms. The most abundant Molecular Function GO terms comprised metal ion binding (GO:0043167; 9,210), DNA binding (GO:0003677; 7,041) and ATP binding (GO:0005524; 6,518). The most represented biological process GO terms were transcription DNA-dependent (GO:0006351; 6,222), signal transduction (GO:0007165; 3,851) and multicellular organismal development (GO: 0007275; 2,908).

**Figure 3.**
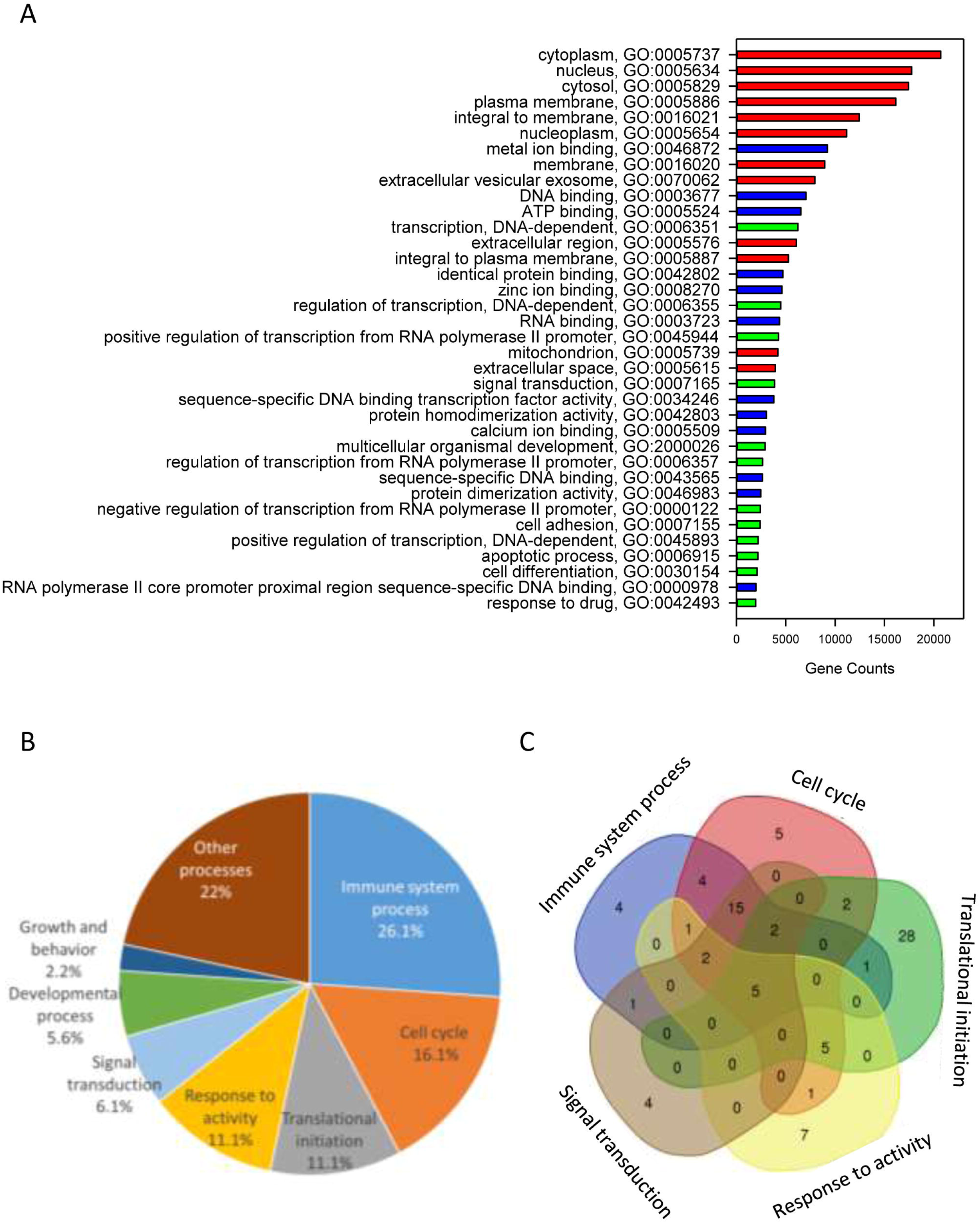
Chimeric genes functional annotation and gene ontology enrichment. **(A)** Gene Ontology (GO) functional annotation analysis over the whole gene model, showing the major GO biological processes (red), GO molecular functions (blue) and GO cellular components (green) for genes found in the gilthead sea bream genome. **(B)** Pie diagram representing the percentage of biological process-enriched GO term functional categories. **(C)** Venn diagram representing the overlapping of the unique gene descriptions between main functional categories.

When tested for enrichment of GO terms among chimeric/composite genes, the 3,648 duplicated genes with 108 non-redundant protein annotations (Supplementary Table 8) rendered 184 enriched biological processes (corrected P-value < 0.05). These genes covering different GO terms related to immune system (26%), cell cycle (16%), translational initiation (11%), response to activity (11%), signal transduction (6%), developmental process (5%) and growth (2%) among others (Figure 3B). The relationship among functional categories is illustrated by a Venn diagram, showing 87 non-redundant gene descriptions of the main five functional categories (Figure 3C). This procedure highlighted that the high representation of immune system in chimeric/composite genes was mostly due to a wide overlapping of immune GO terms with the other enriched functional categories. Intriguingly, main intersections were found among immune system process, cell cycle and signal transduction, comprising 15 enriched GO terms and 15 unique gene descriptions, corresponding to different isoforms of protein NLRC3 and NACTH, LRR and PYD domains-containing protein 12.

### Genome Expansion is Supported by Synteny and Phylogenomic Analyses

Homology relationships between genes contained in the assembled gilthead sea bream super-scaffolds and genes sequenced in other species, as well as their syntenic relationships were studied. From the 30,455 gilthead sea bream genes included in super-scaffolds, 25,806 (84.73%) had orthologs in at least one of the analyzed species, being Nile tilapia (*O. niloticus*, 20,561), zebra mbuna (*M. zebra*, 19,717), platyfish (*X. maculatus*, 15,093) and stickleback (*G. aculeatus,* 14,612) the species sharing more orthologous genes with gilthead sea bream, whereas the lowest numbers of orthologous were obtained in rainbow trout (8,866) and zebrafish (*D. rerio*, 4,288) (Figure 4A). Likewise, the number of syntenic blocks ranged between 483 in *O. niloticus* to 32 in *D. rerio* (Supplementary Table 9). Thus, the levels of both orthology and synteny conservation reflects phylogenetic proximity among the compared species. Also, the number of orthologous genes in syntenic blocks were maximal in *O. niloticus* (9,914; 30.02%), *M. zebra* (9,499; 34.48%) and *G. aculeatus* (6,866; 46.85%), whereas salmonids and cyprinids showed the lowest levels of synteny with 1,284 (*O. mykiss*), 1,482 (Atlantic salmon, *S. salar*) and 44 (*D. rerio*) orthologous in syntenic blocks. The intra-species synteny rendered a total of 268 syntenic blocks in gilthead sea bream that comprised 1,131 paralogs. This feature as well as the high number of connections in the Circos plot of Fig. 4A is indicative of a highly duplicated genome.

**Figure 4.**
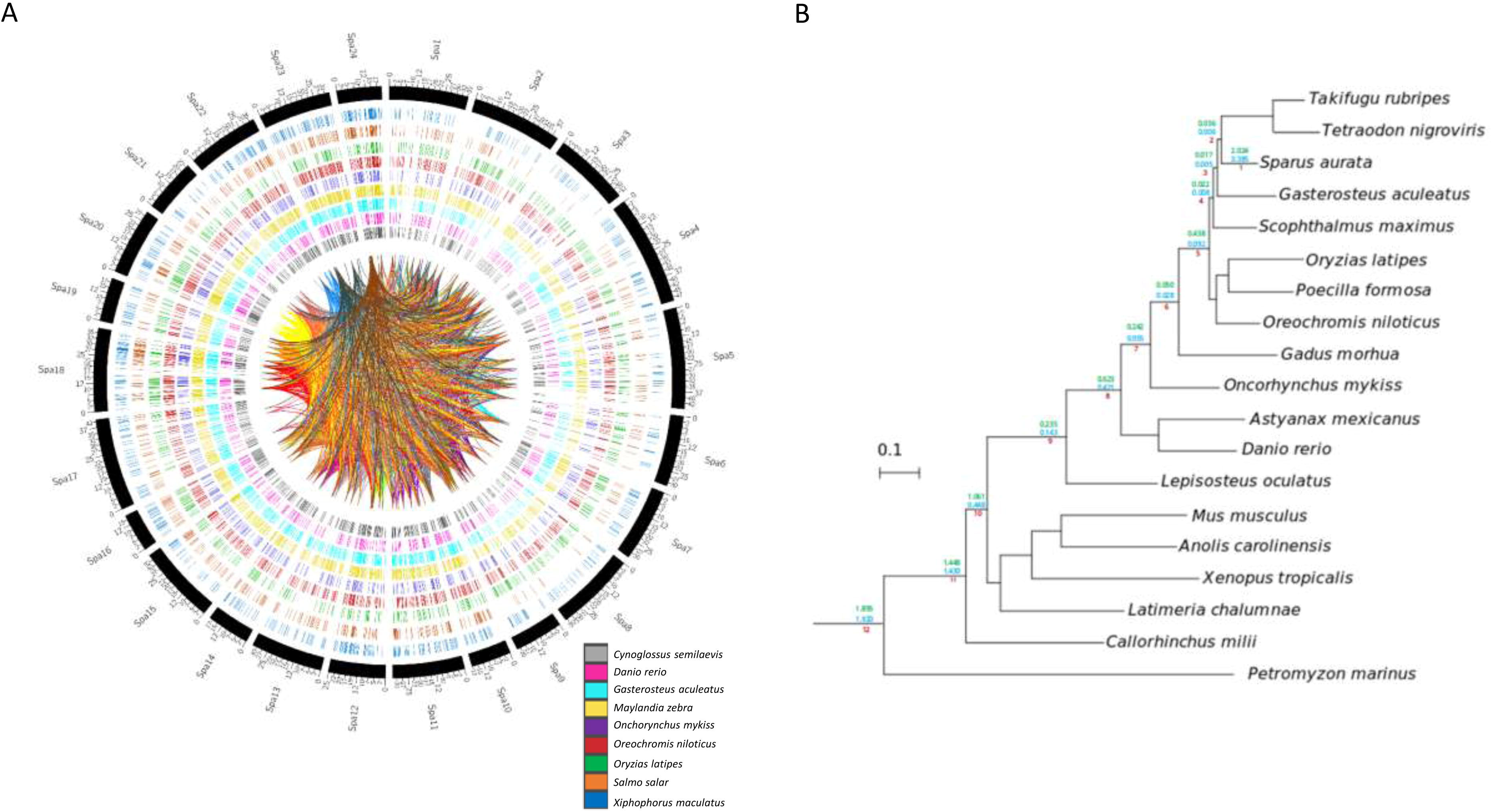
Gene homology and phylogeny of gilthead sea bream. **(A)** Circos plots representing homology relations between gilthead sea bream and other fish species genes. Relations between scaffolded genes with other species with a 99% of identity are shown. Duplicated genes relations between gilthead sea bream chromosomes are represented by inner lines. **(B)** Species tree obtained from the concatenation of 148 single-copy widespread proteins. All nodes are maximally supported (1 aLRT). Number on the branches mark the duplication densities (average number of duplication per gene and per lineage) for gilthead sea bream genes in the lineages leading to this species with (green) or without (blue) expansions.

To gain insights in the evolution of gilthead sea bream genome and study in more detail the origin of these high levels of genomic duplication, we inferred its phylome -i.e. the complete collection of gene evolutionary histories-across nineteen fully-sequenced vertebrate species. To provide a phylogenetic context to our comparisons, we reconstructed a species tree. This was made using two complementary approaches: 1) species tree concatenation of a total of 148 genes with one-to-one orthologous in each of the included species and 2) super-tree reconstruction using 58,484 gene trees from the phylome. Both approaches resulted in the same highly supported topology (Figure 4B), which was fully consistent with the known relationships of the considered species. All trees and alignments are available to browse or download through PhylomeDB (www.phylomedb.org) (Huerta-Cepas et al., 2014) under the phylomeDB ID 714.

From the reconstructed gilthead sea bream phylome, we inferred that 45,162 genes had duplications. The fraction of duplicated genes remained high (17,596) after the removal of gene family expansions (i.e. those resulting in 5 or more in-paralogs). When duplication frequencies per branch in all lineages leading to the gilthead sea bream were computed, two peaks of high duplication ratios (average duplications per gene) were inferred at earliest splits of vertebrates and at the base of teleost fish (teleost-specific genome duplication), which correspond to the known WGDs (Figure 4B; clades 8, 12). Additionally, the gilthead sea bream genome also showed a high rate of species-specific duplications (2.024 duplications per gene; 0.385 duplications per gene after removing expansions). Functional GO enrichment of these duplicated genes highlighted different biological processes, mostly related to genome transposition, immune response and response to stimulus. This referred to the following GO terms: DNA integration (GO:0015074); transposition, DNA-mediated (GO:0006313); RNA-dependent DNA biosynthetic process (GO:0006278); developmental process, (GO:0032502); transposition, RNA-mediated (GO:0032197); DNA recombination, (GO:0006310); immunoglobulin production (GO:0002377); detection of chemical stimulus involved in sensory perception (GO:0050907); regulation of T cell apoptotic process (GO:0070232); telomere maintenance (GO:0000723). In the case of immunoglobulin production, this stated to 24 unique gene descriptions including among others Ig heavy chain Mem5-like isoform X1, Ig heavy chain Mem5-like isoform X2, Ig kappa chain V region 3547, Ig kappa chain V region Mem5, Ig kappa chain V-II region 2S1.3, Ig kappa chain V-IV region Len, Ig lambda chain V-I region BL2, Ig lambda chain V-I region NIG-64, Ig lambda-3 chain C regions, Ig lambda-6 chain C region, Ig lambda-6 chain C region, Ig lambda-like polypeptide 1 isoforms X1, X3 and X4, Ig lambda-like polypeptide 5, pre-B lymphocyte protein 3, integral membrane protein 2A, laminin subunit alpha-2 or Ig kappa chin V19-17. Likewise, the regulation of T cell apoptotic process refers to microfibrillar-associated protein 1, tyrosine-protein kinase JAK2 and JAK3 in addition to different GTPases of IMAP family members (2, 4, 4-like, 8, 8-like). Lastly, the category detection of chemical stimulus involved a wide representation of olfactory receptors, including among others olfactory receptor 10J4-like, 11A11-like, 13C8-like, 146-like, 1M1-like, 2K2-like, 2S2-like, 4C15-like, 4K3-like, 4N5-like, 51G1-like, 5A5-like, 52D1-like, 52K1-like, 5B17-like and 6N1-like.

### Wide Transcriptome Analysis Reveals Different Tissue Gene Duplication Signatures

Up to 70% of the pre-processed reads of the RNA-seq tissue samples were mapped in the assembled genome, yielding 55,423 genes that are reduced to 16,992 after the removal of low expressed genes, low alignments high scoring pairs (HSP) and phylome-based paralogs. From these filtered sequences, up to 5,322 genes were recognized as ubiquitously expressed sequences in the analyzed tissues (Figure 5A). Intestine as a whole (anterior and posterior intestine segments) had the highest number of tissue-exclusive annotated genes (1,198), followed by gills (667), liver (256) and spleen (248) and skeletal white muscle (203). When unique gene descriptions were considered, the order of tissues with a tissue-exclusive number of non-redundant molecular signatures was maintained: intestine (512) > gills (379) > liver (139) > spleen (131) > skeletal muscle (123) (Figure 5B). This yielded a variable percentage of duplicated genes from 28% in the consensus gene list (1.295 out of 4.625) for all the analyzed tissues to 20-17% in muscle and intestine, 12-10% in liver and gills and 6% in spleen. Likewise, the duplication rate ranged between 1.62 from the consensus list to 1.26-1.24 in muscle and intestine, 1.16 in liver, 1.13 in gills and 1.08 in spleen (Figure 5C). The final list of 1,284 tissue-exclusive genes (present in only one tissue) with their number of copies is shown in Supplementary Table 10.

**Figure 5.**
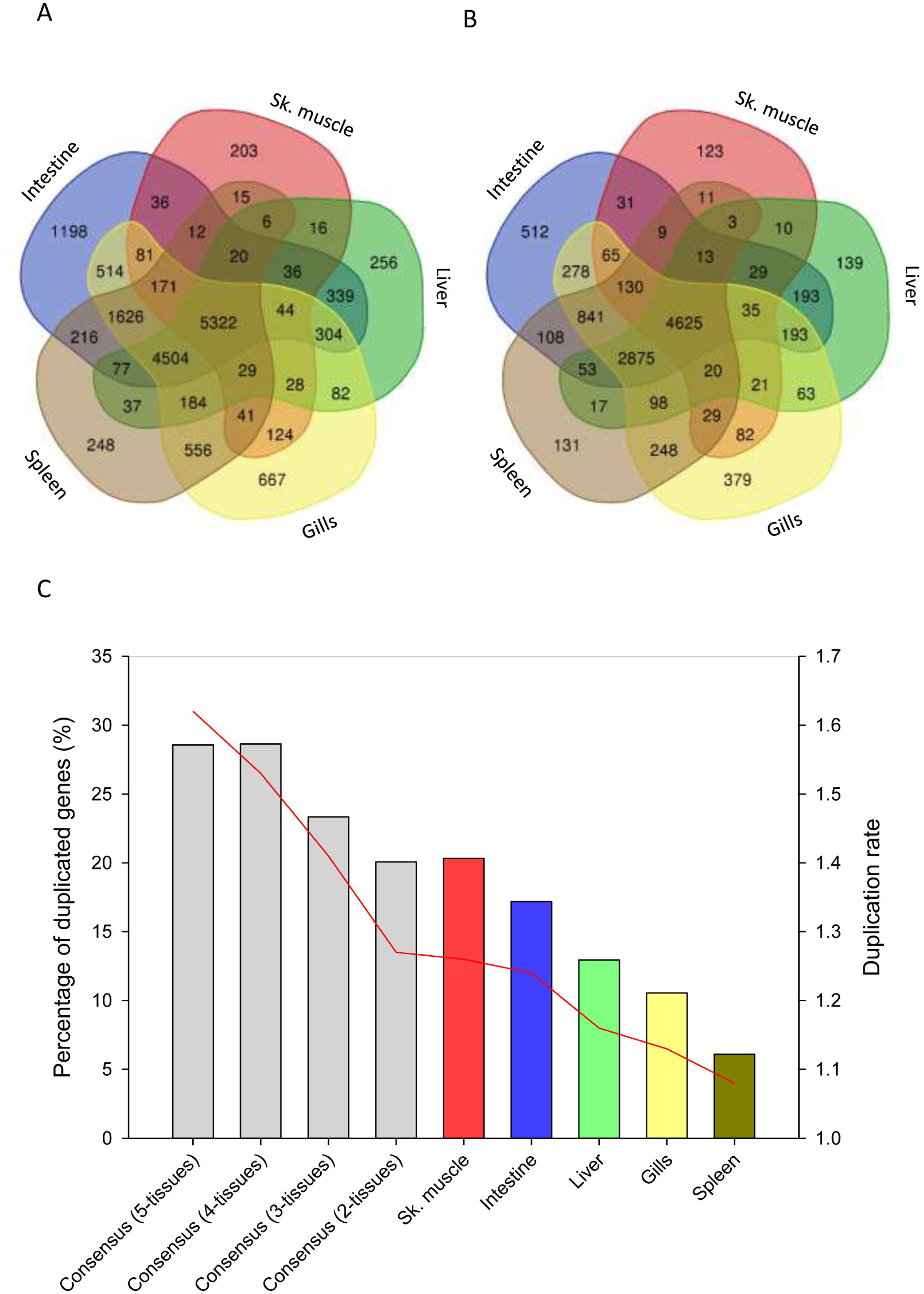
Tissue expression signatures. **(A)** Venn diagram showing the overlap between the gene expression signatures in all analyzed tissues. **(B)** Venn diagram showing the overlap between unique gene annotation expression signatures in all analyzed tissues. Homology-based annotation was done according to the gilthead sea bream transcriptome (Pauletto et al., 2018) and NCBI non-redundant (Nr) database. **(C)** Percentage of duplicated genes among tissues or groups of tissues (blue columns). Red line represents the duplication rate of the unique gene annotations present in a tissue or in a group of tissues.

Tissue-exclusive non-redundant paralogs of intestine, skeletal muscle, liver, spleen and gills are listed in Supplementary Table 11. According to the gene expression pattern in humans and other higher vertebrates (https://www.proteinatlas.org/, https://www.ebi.ac.uk/gxa/home), most of them (65-75%) were classified as tissue- or group-enriched genes (gills paralogs are not included in the analysis due to the lack of a reference expression Atlas for fish species) (Figure 6A). This procedure yielded up to 65 tissue-exclusive paralogs (intestine, 30; skeletal muscle, 17; liver, 13; spleen, 5), showing expression changes between duplicated copies with a similar range of variation when the outliers from intestine (1) and gills (1) were not included in the analysis (Figure 6B). For some of them, including *cav3*, *myod1* and *myod2* (skeletal muscle); *slc6a19* and *aoc1* (intestine); *upp2* and *prom1* (liver); *lmo1* and *yjefn3* (gills); *gp2* and *hbb2* (spleen) the differential gene expression pattern for duplicated genes was validated by qPCR, and overall a high correlation was found for representative genes of all analyzed tissues (Supplementary Table 12).

**Figure 6.**
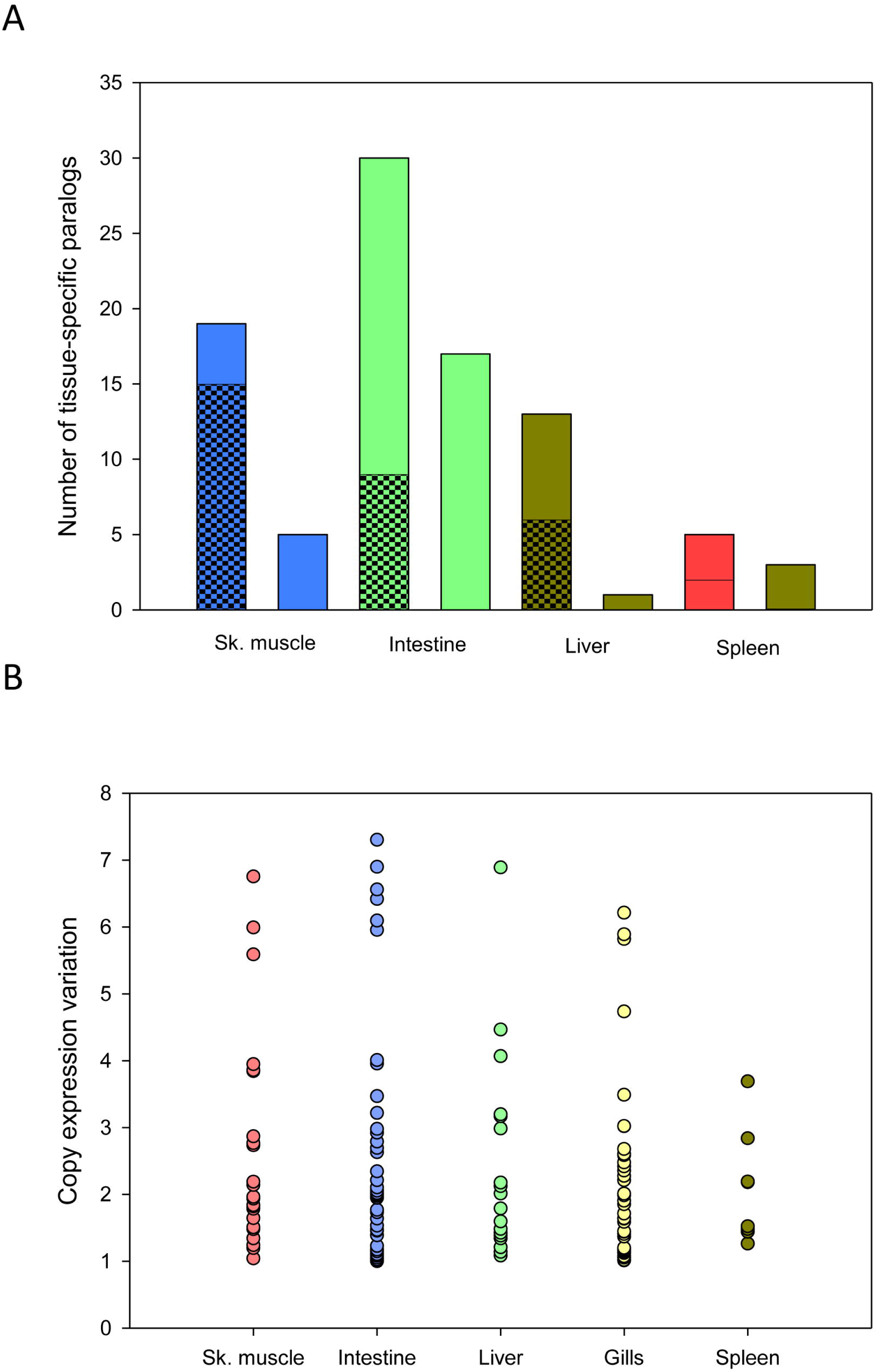
Comparison of tissue-exclusive paralogs and gene expression Atlas in animal models. **(A)** Classification of tissue-exclusive paralog expression enrichment in animal models according to gene expression atlases: enriched in tissue (checkered stacked bar), enriched in tissue and/or other tissues (diagonal stripped stacked bar) and expressed in all tissues (smooth colored column). **(B)** Scatter plot showing the range of expression variation in tissue-exclusive paralogs. Each point represents the variation value for each paralog between the most and the less expressed copies.

## Discussion

Steady advances in sequencing technology and cost reduction are improving the ability to generate high-quality genomic sequences (Metzker, 2010). Certainly, the genome list in the NCBI database (www.ncbi.nlm.nih.gov/genome/browse) contains 340 fish genomes from 248 fish species, with more than 30 corresponding to fish species of special relevance given their economic importance or important role as research model species. In the present study, we have generated and made publicly available a high quality annotated assembly of the gilthead sea bream genome as an effort to generate new genomic tools for a highly cultured fish in all the Mediterranean area. Our sequencing strategy, combining short reads with long read libraries (Nextera MP and PacBio SMRT), has resulted in one of the best fish genome assemblies in terms of number of scaffolds per assembled size (5,039 scaffolds in a 1.24 Gb assembly). Previous attempts in closely related fish resulted in highly fragmented reference genomes due to the use of assembly protocols based solely on short-read sequencing approaches. For instance, the public genomes of European sea bass (*Dicentrarchus labrax*; 680 Mb), spotted green pufferfish (*T. nigroviridis*; 342 Mb) or the Amazon molly (*Poecilia formosa*; 830 Mb) are split in 46,509, 27,918 and 25,474 scaffolds, respectively (Jaillon et al., 2004; Tine et al., 2014; Warren et al., 2018). Likewise, the first gilthead sea bream genome draft comprised 55,202 scaffolds in a 760 Mb assembly (Pauletto et al., 2018). In concurrence with the present study, a new genome draft of gilthead sea bream was submitted to NCBI (Bioproject accession PRJEB31901), comprising ∼833 Mb, which is still below our assembly. This yielded a higher number of unique gene annotated descriptions when comparing our assembled genome with the two previous releases (21,275 *vs.* 13,835-19,631).

Fish comprise the largest and most diverse group of vertebrates, ranging the size of sequenced genomes between 342 Mb in *T. nigroviridis* to 2.90 Gb in *S. salar* (Yuan et al., 2018). Our unmasked assembled genome is, thereby, of intermediate size (1.24 Gb), although the full genome is expected to be around 350 Mb longer. Indeed, the current assembly contains more than 5,000 unique gene descriptions that are not present in the super-scaffolding based on the first genome draft (Pauletto et al., 2018). Estimations of gilthead sea bream genome size based on flow cytometry of red blood cells rendered a smaller genome size (∼930 Mb) (Peruzzi et al., 2005). Nevertheless, the accuracy of the technique is limited due to high intra- (up to 10%) and inter-assay (20-26%) sources of variation (Pedersen, 1971; Gregory, 2005). Certainly, differences in internal/external genome size standards, sample preparation, staining strategies or stochastic drift of instruments might result in significant differences in such genome size estimations (Doležel et al., 1998), and consequently computational methods (e.g. k-mer frequency counts) are emerging as more reliable approaches for genome size estimations (Sun et al., 2018).

Another important output from our k-mer count analysis was a pronounced second peak that is indicative of a high amount of repeated sequences. In this regard, the results of redundancy analysis based on actively transcribed genes approximated a low fraction of segmental duplications (1.01%) that is indicative of a reduced genome mis-assembly (Kelley and Salzberg, 2010). Accordingly, most of the gene predictions reported by us showed a sufficient degree of divergence to support the idea of true gene expansions. Reliable gene duplication was also supported by synteny analysis, which makes difficult to establish inter-species synteny blocks probably as the result of the over-representation of gene expansions during the recent evolution of the gilthead sea bream lineage. This was confirmed by phylome analysis, which showed an average of 2.024 copies for the 55,423 actively transcribed genes, in at least one of the analyzed tissues as a representation of metabolically- and immune-relevant tissues. This number of tissue-regulated transcripts with a high percentage of duplications offers the possibility of an enhanced adaptive plasticity in a challenging evolutionary environment. Certainly, paralog retention in fish is usually related to specific adaptive traits driven by their particular environments (Maere et al., 2005). Examples of this are the expansion of the antifreeze glycoprotein Afgp in Antartic notothenioid fish (Chen et al., 2008) or the claudins and aquaporins in European sea bass (Tine et al., 2014). At the global level, the highest percentages of duplicated genes are reported for eel (36.6%) and zebrafish (31.9%) (Inoue et al., 2015), but intriguingly the values reported by us in gilthead sea bream (56.5%) are even higher for the duplication ratio calculated as the percentage of non-redundant duplicated annotations.

Importantly, gene functional enrichment in lineage-specific duplicated genes of gilthead sea bream evidenced an increased presence of DNA integration, transposition and immunoglobulin production. This finding suggests that most of the expansions undergone by the gilthead sea bream genome derive from the activities of MGEs and from the immune response as key processes in the species adaptability. Immune genes play a crucial role in the survival and environmental adaptation of species, and are particularly important in aquatic animals, which are continuously and directly exposed to an environment with water-borne pathogens. Thus, duplicate retentions and tandem repeats are commonly found among fish immune genes, with special relevance in those involved in pathogen recognition systems and inhibitors/activators of inflammation (Howe et al., 2016; Li et al., 2017). In fact, the immunoglobulin loci of teleosts are among the largest and most complex described, sometimes containing even several hundreds of V genes (Fillatreau et al., 2013). This scenario seems to be likely orchestrated by selfish elements (introns, repeats, transposons, gene families), which trigger genomic rearrangements, substitutions, deletions and insertions (Kidwell, 2002), leading to the increment of size and complexity of the genome in addition to new gene combinations that result in modified or new biological functions (Lynch and Conery, 2000).

The characterized mobilome highlighted an abundant representation of MGEs as well as a number of chimeric genes that apparently evolved from the co-domestication and/or co-option of MGEs. Co-option is indeed a recurrent mechanism that has contributed to innovations at various levels of cell signalling and gene expression several times during the evolution of vertebrates (Arkhipova et al., 2012). The most represented source of gene co-option in our gilthead sea bream genome were LINE retrotransposons and *Tc1/Mariner* DNA-transposons, which have been extensively reported in mammalian models as examples of transposable elements domestication (Jangam et al., 2017). Among these chimeric genes (Supplementary Table 7), a relevant number of NOD-like receptors (NLRs), including NACHT-, LRR- and PYD-containing proteins (NLRP) and NOD-like receptor CARD domains (NLRCs), emerged. These receptors are innate sensors involved in intracellular monitoring to detect pathogens that have escaped to extracellular and endosomal surveillance. Fish are in fact the first in evolution to possess a fully developed adaptive immune system. However, due to the environment they live in, they still rely on and maintain a wide array of innate effectors, showing an impressive species-specific expansion of these genes (Stein et al., 2007), as is the case for the more than 400 NLR family members in zebrafish (Li et al., 2017). These duplications reflect the evolutionary need of detecting threats in a pathogen rich environment, and correlate to the diversity of habitats with species-specific traits in teleosts, the largest group of vertebrates.

Analysis of RNA-seq active transcripts across five different tissues also pointed out the association of gene duplication with different tissue expression patterns. Indeed, gene duplication and subsequent divergence is basic for the evolution of gene functions, although the role of positive selection in the fixation of duplicated genes remains an open question (Kidwell, 2002; Kondrashov, 2012). A highly conservative filtering step was applied in our gene dataset in order to avoid genetic redundancy or pseudogeneization that could be potentially mistaken as true duplication events (Innan and Kondrashov, 2010). This procedure showed higher duplication levels in genes expressed in two or more tissues as compared to those with a tissue-exclusive expression, being in accordance the annotation and functions of the tissue-exclusive paralogs with the reference Atlas of tissue gene expression of higher vertebrates. This fact is in agreement with earlier studies demonstrating that in a tissue functionalization context (i.e. gene copies expressed in several tissues), gene duplication leads to increased levels of tissue specificity (Huerta-Cepas et al., 2011). Likewise, we observed herein that gene copies expressed in two or more tissues showed increased duplication rates and percentages of retained paralogs in comparison to tissue-exclusive genes. Analysis of qPCR, designed to discriminate the expression patterns of selected tissue-exclusive paralogs (liver, 2; skeletal muscle, 3; intestine, 2; gills, 2; spleen, 2), further emphasized this functional divergence towards a more specific regulation of duplicated genes. However, future studies (combining both targeted and untargeted transcriptome approaches) are still needed to clarify the relationship between the gene expressions of duplicated genes and specific phenotypic traits. Although at this stage, it appears conclusive that the genome of gilthead sea bream has retained an increased number of duplications in comparison to closest relatives. In comparison to other modern fish lineages, this higher gene duplication ratio is also extensive to salmonids and cyprinids (Macqueen and Johnston, 2014; Chen et al., 2019) that still conserved signatures of a WGD in their genome. Since the gene repertory of gilthead sea bream is also characterized by the persistence of multiple gene copies for a given duplication, it is likely that this feature is mostly the result of highly active MGEs, allowing the improved plasticity across the evolution of a fish family with a remarkable habitat diversification (Sbragaglia et al., 2019). This observation, together with a recent eel transcriptome study, renew the discussion about fish lineage specific re-diploidization after 3R or even an additional WGD (Rozenfeld et al., 2019).

In summary, a combined sequencing strategy of short- and long-reads produced a high quality draft of gilthead sea bream genome that can be accessed by a specific genome browser that includes a karyotype alignment. The high coverage and depth of this assembly result in a valuable resource for forthcoming NGS-based applications (such as RNA-seq or Methyl-seq), metatranscriptome analysis, quantitative trait loci (QTLs) and gene spatial organization studies conducted to improve the traits of this highly cultured farmed fish. Assembly analysis suggests that transposable elements are probably the major cause of the enlarged genome size with a high number of functionally specialized paralogs under tissue-exclusive regulation. These findings highlight the genome plasticity of a protandric, euryhalin and eurytherm fish species, offering the possibility to further orientate domestication and selective breeding towards more robust and efficient fish, making gilthead sea bream an excellent model to investigate the processes driving genome expansion in higher vertebrates.

### Data Availability

Raw sequence reads generated during the current study were deposited in the Sequence Read Archive of the National Center for Biotechnology Information (NCBI). Primary accession numbers: PRJNA551969 (Bioproject ID); SAMN12172390-SAMN12172427 (genomic Illumina Nextseq500 PE, MP and PacBio RS II raw reads); SAMN12172428-SAMN12172433 (RNA-seq Illumina NextSeq500 SE raw reads from skeletal white muscle); SAMN12172434, SAMN12172435 (RNA-seq Illumina NextSeq500 PE raw reads from anterior and posterior intestine). PRJNA507368 (Bioproject ID for raw reads from gills, liver and spleen tissues); SRR8255950, SRR8255962-70 (RNA-seq Illumina NextSeq500 raw reads from gills, liver and spleen tissues). All phylogenetic trees and alignments of the gilthead sea bream genome are publicly available through phylomeDB (http://www.phylomedb.org, phylome ID 714). A genome browser was built for the navigation and query of the assembled sequences in http://nutrigroup-iats.org/seabreambrowser.

## Conflict of Interest

The authors declare that the research was conducted in the absence of any commercial or financial relationships that could be construed as a potential conflict of interest.

## Author contributions

This study was designed and coordinated by JP-S. Material from gilthead sea bream used for genome sequencing was extracted by J-AC-G and JP-S. Genome assembly and annotation were performed by BS and CL. Evolutionary and phylogenomics analysis were performed by TG. Genome browser was implemented by AH. Data analysis and integration were performed by FN-C, J-AC-G, M-CP, AS-B and JP-S. All authors read, discussed, edited and approved the final manuscript.

## Funding

This work was financed by Spanish (Intramural CSIC, 1201530E025; MICINN BreamAquaINTECH, RTI2018-094128-B-I00) and European Union (AQUAEXCEL^2020^, 652831) projects to JP-S. BS was supported by a predoctoral research fellowship (Doctorados industriales, DI-17-09134) from Spanish MINECO.

## Acknowledgements

The authors thank M. A. González for technical assistance with gene expression analyses.

## Supplementary Figures

**Supplementary Figure 1. K-mer based genome estimation size and scaffold distribution. (A)** 63-mer frequency histogram for the gilthead sea bream assembly for genome size estimation. **(B)** Cumulative length of the assembled scaffolds fitted to total scaffold length. Highlighted points remark the number of scaffolds compressed under 25, 50, 75 and 90% of the total scaffold length.

**Supplementary Figure 2. Reconstructed gilthead sea bream super-scaffolds**. All scaffolds (1.87-12.05 Mb) were anchored to the gilthead sea bream chromosomes (2*n=48*). Scaffolds are listed at the right side of each super-scaffold, and a nucleotide position of reference for the browser is marked in the left side. A genome browser to access and navigate the super-scaffold is available at http://nutrigroup.iats.org/seabreambrowser.

**Supplementary Figure 3. MGEs and chimeric genes KRONA representation.** KRONA representation of the distribution of all MGEs and chimeric genes belonging to the mobilome draft of the gilthead sea bream excluding low complexity repeats and introns.

## Supplementary Tables

Supplementary Table 1. Forward and reverse primers used for real-time qPCR.

Supplementary Table 2. Summary statistics of sequencing data, detailed for each sequencing strategy.

**Supplementary Table 3. Assembly metrics for the gilthead sea bream genome**. Metrics were inferred using the script assemblathon_stats.pl available at http://korflab.ucdavis.edu/datasets/Assemblathon/Assemblathon2/Basic_metrics/assemblathon_stats.pl.

**Supplementary Table 4. Dedupe redundancy analysis with nucleotide sequences**. Analysis was performed over the nucleotide sequences of the final set of active transcripts retrieved from RNA-seq transcriptome analysis.

Supplementary Table 5. MGEs and chimeric related-genes found in the mobilome draft of gilthead sea bream genome.

Supplementary Table 6. Predicted and annotated non coding RNAs in the gilthead sea bream genome.

Supplementary Table 7. Summary of annotations of chimeric/composite genes and multigene families of the gilthead sea bream genome including BLAST hits and statistics of those presenting homology to MGEs.

**Supplementary Table 8. Biological process GO term enrichment results in transposon-overlapping gene fraction.** Supplementary Table shows the GO annotation of the 108 non-redundant descriptions corresponding to chimeric/composite genes.

Supplementary Table 9. Synteny results between gilthead sea bream and related species.

**Supplementary Table 10. Tissue-exclusive genes dataset.** Homology-based annotation was done according to the gilthead sea bream transcriptome and NCBI non-redundant (Nr) database, and the correspondent Uniprot KB AC/ID was retrieved for each gene. The number of copies is shown in Copy number column.

**Supplementary Table 11. Tissue-exclusive duplicated gene list.** Results highlights tissue-expression pattern in other animal models: enriched in tissue (red), enriched in tissue and/or other tissues (green), expressed in all tissue (blue) and unclassified (uncolored). A range of colors is shown for the Δ_copies_ between paralog sets ordered by each category. Column Corrected P-val shows the result for the ANOVA (FDR < 0.05) test.

**Supplementary Table 12. Pearson correlation coefficients between RNA-seq and real-time qPCR expression values of tissue-exclusive genes**. AI-PI: Anterior & Posterior intestine; WSM: White skeletal muscle; L: Liver; S: Spleen; G: Gills. PCC: Pearson correlation coefficient. ^1^P-value obtained in Pearson correlation.

