## Supplementary Table 1 for "Genome Sequencing and Transcriptome Analysis Reveal Recent Species-specific Gene Duplications in the Plastic Gilthead Sea Bream"

| **Supplementary Table 1**. Forward and reverse primers used for real-time qPCR. | | |
| --- | --- | --- |
| **Gene name** | **Symbol** | **Primer sequence** |
| Amiloride-sensitive amine oxidase [copper-containing] (copy a) | *aoc1a* | F: TGC TGG TTG GCT TGG CTG GTT |
|  |  | R: GCA CCA TGA TGA GCC CAC TCT CTT GA |
| Amiloride-sensitive amine oxidase [copper-containing] (copy b) | *aoc1b* | F: CCT ACC GAG TCC GCT CAT CAG |
|  |  | R: GTG TCA CCA GCA TAG AAG GCA ATA GC |
| Caveolin 3 (copy a) | *cav3a* | F: GAT CAG TAC CAA TAC ACC AAC |
|  |  | R: GTA GCA CCA GTA CTT GGA |
| Caveolin 3 (copy b) | *cav3b* | F: ACT GTG TCC AGT ATG GTG CTA C |
|  |  | R: GGA AGC CCC AAG AGC AGA G |
| Cadherin-15 | *cdh15* | F: AAC GCT TAT CTG AGC TAC TCT ATC ATT GG |
|  |  | R: CTG GTT GTT GAT ACC GAA CAT TGT CTT G |
| Claudin-15 | *cldn15* | F: CCG ATT GTG GAA GTA GTG GCT CTG GT |
|  |  | R: CAG CAT CAC CCA ACC GAC GAA CC |
| C-type lectin domain family 10 member A | *clec10a* | F: CGA CTC TGG ACT CCC TCA |
|  |  | R: CGT TGT TGA TGG TGC GTT C |
| pancreatic secretory granule membrane major glycoprotein GP2-like (copy a) | *gp2a* | F: GCT GCC CGG TGT TGT CAA AAC AGA |
|  |  | R: CCC TCG CCG TTC TTC AGC |
| pancreatic secretory granule membrane major glycoprotein GP2-like (copy b) | *gp2b* | F: CCA GTG AAG ATG GAT TGA TGA GGA C |
|  |  | R: GGA CTC CGC CTG CAT TCG |
| Uridine phosphorylase 2 (copy a) | *upp2a* | F: CGA GGG AAC TGA TCG CTA CTG |
|  |  | R: GTG CAG CAT GAA TGG AGA TGG AA |
| Uridine phosphorylase 2 (copy b) | *upp2b* | F: ACT GAG CTG GAC GAG GGA |
|  |  | R: TCC GAT CAC TGT GGG AAT GT |
| YjeF N-terminal domain-containing protein 3 (copy a) | *yjefn3a* | F: AGA GCT ACT GAG GGA CTA |
|  |  | R: AGA TGA CTA GCA CTG TTG |
| YjeF N-terminal domain-containing protein 3 (copy b) | *yjefn3b* | F: AGG TTC TCG GGG AAG CAT TTC TT |
|  |  | R: AAT TCT ATG AGA CAG TCT GTG CCA GGA TA |
| Hemoglobin subunit beta-2 (copy a) | *hbb2a* | F: GTG GAA CAA ATC TAC CGG CAA A |
|  |  | R: AAC GAC CTC ATA CTT CAT CTT GGA |
| Hemoglobin subunit beta-2 (copy b) | *hbb2b* | F: GTG GAA CAA ATC TAC CGG CAA A |
|  |  | R: AAC GAC CTC ATA CTT CAT CTT GGA |
| Rhombotin-1 (copy a) | *lmo1a* | F: CTC AAG AAC AAC ATG ATC CTG TGC CA |
|  |  | R: GCC TCT CTG CGC TGC CAT TA |
| Rhombotin-1 (copy b) | *lmo1b* | F: CCA TTT AGA CTG CTT CGC CTG T |
|  |  | R: AGA TGG CCT CCT TCA TAG TCC |
| Galectin-1 | *lgals1* | F: GTG TGA GGA GGT CCG TGA TG |
|  |  | R: ACT GTA GAG CCG TCC GAT AGG |
| Myogenic differentiation factor 1 (copy a) | *myod1a* | F: GCG GCT GGT CCA TGT GGG |
|  |  | R: GGA GGG TGA AAC TGA AGA GGA GGA |
| Myogenic differentiation factor 1 (copy b) | *myod1b* | F: CTG TTT CAC CCT CCC TTC CT |
|  |  | R: AGG TGC AGC AGG GAT GAC |
| Myogenic differentiation factor 2 (copy a) | *myod2a* | F: CTC TCC GTG TTC GAG CAC AAG TGA |
|  |  | R: TGC TTC TCC TGG ACG TAT GCT GTC T |
| Myogenic differentiation factor 2 (copy b) | *myod2b* | F: GCA ACC AGA CAG CAT ACG AGT CCA |
|  |  | R: CTG TGC TGA TCC GCT CTA CGA TG |
| prominin-1 isoform X4 (copy a) | *prom1a* | F: GCA CCA GAG ACA GAG GAA GAA CT |
|  |  | R: CGA TGA AGA TGG AGG TAG CGA TGA |
| prominin-1 isoform X4 (copy b) | *prom1b* | F: GGT CAA GGT CAC GAA TAT CCT GAG T |
|  |  | R: CAC CAC ATG CGA GGC GTT ATG |
| sodium-dependent neutral amino acid transporter B(0)AT1-like (copy a) | *slc6a19a* | F: GTG GGC TTC TGT GTG GGA CTT |
|  |  | R: CTC CTC CGT GGC TCT GAC A |
| sodium-dependent neutral amino acid transporter B(0)AT1-like (copy b) | *slc6a19b* | F: AAT GCC TCT GCT GCT GCT |
|  |  | R: TCC CAC ACT GCC CTT CCT |
| Transcription factor SOX3 | *sox3* | F: ACA TGA AGG AGC ACC CGG ATT ATA AAT ACC |
|  |  | R: GGG CAA AGA ATA CTT GTC TTT CTT GAG CAA |
