## Supplementary material for "Genome Sequencing and Transcriptome Analysis Reveal Recent Species-specific Gene Duplications in the Plastic Gilthead Sea Bream": Supplentary Table 2

| **Supplementary Table 2**. Summary statistics of sequencing data, detailed for each sequencing strategy. | | | | | |
| --- | --- | --- | --- | --- | --- |
| **Platform** | **Illumina Nextseq500** | | | | **PacBio RS II** |
| Read type (insert size) | Paired-End (360 bp) | Paired-End (747 bp) | Mate-Pair (5kb) | Mate-pair (8kb) | smRT (<50kb) |
| Read length (bp) | 2x150 | 2x150 | 2x75 | 2x75 | 75-49000 |
| Read number | 2x3.27E+08 | 2x3.05E+08 | 2x7.17E+07 | 2x6.38E+07 | 1.02E+06 |
| Total length (bp) | 2x4.57E+10 | 2x4.91E+10 | 2x5.38E+09 | 2x6.38E+09 | 8.55E+09 |
| Sequencing depth | 37x | 39x | 4.5x | 4x | 7x |
