## Supplementary Table 3 for "Genome Sequencing and Transcriptome Analysis Reveal Recent Species-specific Gene Duplications in the Plastic Gilthead Sea Bream"

| **Supplementary Table 3.** Assembly metrics of sea bream genome. | |
| --- | --- |
| Number of scaffolds | 5,039 |
| Total size of scaffolds | 1,246,531,774 |
| Longest scaffold | 16,075,163 |
| Shortest scaffold | 765 |
| Number of scaffolds > 1K nt | 5,037 (99.96%) |
| Number of scaffolds > 10K nt | 3,724 (73.90%) |
| Number of scaffolds > 100K nt | 1,977 (39.23%) |
| Number of scaffolds > 1M nt | 244 (4.84%) |
| Number of scaffolds > 10M nt | 4 (0.08%) |
| Mean scaffold size | 247,376 |
| Median scaffold size | 56,683 |
| N50 scaffold length | 1,073,340 |
| L50 scaffold count | 227 |
| Scaffold %A | 27.66 |
| Scaffold %C | 19.9 |
| Scaffold %G | 19.92 |
| Scaffold %T | 27.63 |
| Scaffold %N | 4.89 |
| Scaffold %non-ACGTN | 0 |
| Number of scaffold non-ACGTN nt | 0 |
| Percentage of assembly in scaffolded contigs | 99.2% |
| Percentage of assembly in unscaffolded contigs | 0.8% |
| Average number of contigs per scaffold | 10.3 |
| Average length of break (>25 Ns) between contigs in scaffold | 1,297 |
| Number of contigs | 51,918 |
| Number of contigs in scaffolds | 50,568 |
| Number of contigs not in scaffolds | 1,350 |
| Total size of contigs | 1,185,707,601 |
| Longest contig | 538,223 |
| Shortest contig | 3 |
| Number of contigs > 1K nt | 49,226 (94.8%) |
| Number of contigs > 10K nt | 25,806 (49.7%) |
| Number of contigs > 100K nt | 1960 (3.8%) |
| Number of contigs > 1M nt | 0 (0.00%) |
| Number of contigs > 10M nt | 0 (0.00%) |
| Mean contig size | 22,838 |
| Median contig size | 9,883 |
| N50 contig length | 53,619 |
| L50 contig count | 6,237 |
| Contig %A | 29.08 |
| Contig %C | 20.93 |
| Contig %G | 20.94 |
| Contig %T | 29.05 |
| Contig %N | 0.01 |
| Contig %non-ACGTN | 0 |
| Number of contig non-ACGTN nt | 0 |
