## Supplementary Table 4 for "Genome Sequencing and Transcriptome Analysis Reveal Recent Species-specific Gene Duplications in the Plastic Gilthead Sea Bream"

| **Supplementary Table 4**. **Dedupe redundancy analysis with nucleotide sequences.** Analysis was performed over the nucleotide sequences of the final set of active transcripts retrieved from RNA-seq transcriptome analysis. | | | | | | |
| --- | --- | --- | --- | --- | --- | --- |
|  | **98% threshold** | | **95% threshold** | | **90% threshold** | |
| **Input protein sequences** | 54,423 | 100% | 54,423 | 100% | 54,423 | 100% |
| **Duplicates** | 559 | 1.01% | 559 | 1.01% | 559 | 1.01% |
| **Containments** | 1,836 | 3.31% | 2,800 | 5.05% | 3,786 | 6.83% |
| **Clusters** | 53,028 | 95.68% | 52,064 | 93.94% | 51,078 | 92.16% |
