## Supplementary Table 5 for "Genome Sequencing and Transcriptome Analysis Reveal Recent Species-specific Gene Duplications in the Plastic Gilthead Sea Bream"

| **Supplementary Table 4**. Summary of MGEs and chimeric related-genes found in the mobilome draft of gilthead sea bream genome. | | | | | |
| --- | --- | --- | --- | --- | --- |
| **Class** | **Type** | **MGE**  **Groups** | **Size (bp)**  **in genome** | **Percentage**  **of mobilome** | **Percentage**  **of genome** |
| Class I | LTR retroelements | 4 | 27,226,719 | 2.88 | 2.18 |
|  | non-LTR retrotransposons | 14 | 27,743,197 | 2.94 | 2.23 |
|  | YR retrotransposons | 1 | 222,290 | 0.02 | 0.02 |
| Class II | DNA transposons | 27 | 99,635,002 | 10.55 | 7.99 |
| Introns | Introns | 1 | 598,945,346 | 63.44 | 48.05 |
| Chimeric/composite  Multicopy genes | Non-LTR retrotransposons-related genes | 3 | 7,390,894 | 0.78 | 0.59 |
|  | LTR retroelements-related genes | 4 | 729,596 | 0.08 | 0.06 |
|  | DNA transposons-related genes | 10 | 5,867,940 | 0.62 | 0.47 |
|  | unknown | 2 | 4,107,023 | 0.44 | 0.33 |
|  | ncRNA genes | 1 | 53,849 | 0.01 | 0.004 |
|  | YR retroelement-related genes | 1 | 23,583 | 0.002 | 0.002 |
|  | Genes with repeats | 1 | 1,630 | 0.0002 | 0.0001 |
|  | Viral-type genes | 1 | 214,839 | 0.023 | 0.017 |
|  | Clan AA peptidase genes | 11 | 47,283 | 0.005 | 0.004 |
|  | Scan/Krab genes | 1 | 2,801 | 0.0003 | 0.0002 |
| ncRNA  genes | Long ncRNA genes | 10 | 10,750,971 | 1.14 | 0.86 |
|  | Small ncRNA genes | 11 | 1,036,782 | 0.11 | 0.083 |
| Repetitive  DNA | De novo repeats | 2500 | 159,623,699 | 16.91 | 12.81 |
|  | Known repeats | 5 | 475,972 | 0.05 | 0.04 |
| Total Mobilome size | |  | 944,099,416 | 100 | 75.7 |
| Fraction without including introns | |  | 345,154,070 | 36.5 | 27.6 |
