## Supplementary Table 6 for "Genome Sequencing and Transcriptome Analysis Reveal Recent Species-specific Gene Duplications in the Plastic Gilthead Sea Bream"

| **Supplementary Table 5**: non-coding RNAs predicted and annotated in the sea bream genome. | | | | |
| --- | --- | --- | --- | --- |
|  | **Type of ncRNA** | **ncRNA Subtypes** | **ncRNAs with gene name** | **ncRNAs annotated** |
| lncRNA | Processed_pseudogene | 16 | 13 | 100 |
|  | Processed_transcript | 45 | 151 | 151 |
|  | Pseudogene | 178 | 3 | 942 |
|  | Antisense | 37 | 136 | 136 |
|  | IG_V_pseudogene | 1 | 1 | 1 |
|  | TR_V_gene | 5 | 15 | 15 |
|  | lincRNA | 1,089 | 134 | 5,528 |
|  | Sense_intronic | 5 | 35 | 35 |
|  | Misc_RNA | 17 | 21 | 22 |
|  | Unprocessed_pseudogene | 25 | 45 | 45 |
|  | Subtotal lncRNAs | 1,418 | 554 | 6,975 |
| sncRNA | miRNA | 806 | 1,166 | 5,668 |
|  | rRNA | 69 | 92 | 104 |
|  | snoRNA | 221 | 288 | 496 |
|  | snRNA | 144 | 183 | 237 |
|  | sRNA | 4 | 6 | 6 |
|  | tRNA | 329 | 0 | 2,272 |
|  | Subtotal sncRNAs | 1,573 | 1,735 | 8,783 |
| **Total ncRNAs (lncRNAs and sncRNAs)** | | **2,991** | **2,289** | **15,758** |
