## Supplementary Table 9 for "Genome Sequencing and Transcriptome Analysis Reveal Recent Species-specific Gene Duplications in the Plastic Gilthead Sea Bream"

| **Supplementary Table S6**. Synteny between sea bream and related species | | | | | | | |
| --- | --- | --- | --- | --- | --- | --- | --- |
| **Species** | **Homologous genes in species** | **Syntenic**  **blocks** | **Number of scaffolds**  **(sea bream)** | **Syntenic genes in Sparus aurata (%)** | **Number of chromosomes (species)** | **Syntenic genes in Species (%)** | **Diploid number** |
| *Gasterosteus aculeatus* | 14,655 | 381 | 271 | 10.68 | 21 | 46.85 | 2n=42 |
| *Maylandia zebra* | 27,550 | 446 | 344 | 14.35 | 22 | 34.48 | 2n=44 |
| *Oreochromis niloticus* | 33,024 | 483 | 364 | 15.73 | 23 | 30.02 | 2n=44 |
| *Xiphophorus maculatus* | 21,989 | 385 | 299 | 11.67 | 24 | 29.71 | 2n=48 |
| *Oryzias latipes* | 21,031 | 347 | 250 | 10.11 | 24 | 25.04 | 2n=48 |
| *Cynoglossus semilaevis* | 17,797 | 386 | 210 | 8.00 | 21 | 24.91 | 2n=42 |
| *Oncorhynchus mykiss* | 19,396 | 245 | 114 | 4.01 | 29 | 6.62 | 2n=58 |
| *Salmo salar* | 27,693 | 271 | 116 | 4.35 | 28 | 5.35 | 2n=58 |
| *Sparus aurata* | 55,423 | 268 | 100 | 1.45 | 24 | 2.04 | 2n=48 |
| *Danio rerio* | 7,564 | 32 | 17 | 0.25 | 11 | 0.58 | 2n=50 |
