## Supplementary Table 12 for "Genome Sequencing and Transcriptome Analysis Reveal Recent Species-specific Gene Duplications in the Plastic Gilthead Sea Bream"

| **Supplementary Table 9**. Pearson correlation coefficients between RNA-seq and real-time qPCR expression values of tissue-exclusive genes. | | | |
| --- | --- | --- | --- |
| **Gene** | **Tissue** | **PCC** | **p^1^** |
| *aoc1a* | AI-PI | 0.905 | 0.090 |
| *aoc1b* | AI-PI | 0.630 | 0.370 |
| *cav3a* | WSM | 0.991 | <0.001 |
| *cav3b* | WSM | 0.773 | 0.070 |
| *cdh15* | WSM | 0.702 | 0.120 |
| *cldn15* | AI-PI | 0.884 | 0.116 |
| *clec10a* | L | 0.971 | 0.029 |
| *gp2a* | S | 1.000 | 0.010 |
| *gp2b* | S | 0.928 | 0.244 |
| *hbb2a* | S | 0.999 | 0.030 |
| *hbb2b* | S | 0.979 | 0.129 |
| *Imo1a* | G | 0.998 | 0.030 |
| *Imo1b* | G | 1.000 | 0.007 |
| *lgals1* | S | 0.715 | 0.493 |
| *myod1a* | WSM | 0.540 | 0.269 |
| *myod1b* | WSM | 0.653 | 0.159 |
| *myod2a* | WSM | 0.791 | 0.060 |
| *myod2b* | WSM | 0.402 | 0.430 |
| *prom1a* | L | 0.977 | 0.023 |
| *prom1b* | L | 0.994 | 0.006 |
| *slc6a19a* | AI-PI | 0.740 | 0.260 |
| *slc6a19b* | AI-PI | 0.920 | 0.080 |
| *sox3* | G | 1.000 | 0.010 |
| *upp2a* | L | 0.986 | 0.014 |
| *upp2b* | L | 0.956 | 0.040 |
| *yjefn3a* | G | 0.999 | 0.020 |
| *yjefn3b* | G | 0.932 | 0.236 |
| AI-PI: Anterior & Posterior intestine; WSM: White skeletal muscle; L: Liver; S: Spleen; G: Gills. PCC: Pearson correlation coefficient. ^1^P-value obtained in Pearson correlation. | | | |
