## Supplementary figures and images for "Genome Sequencing and Transcriptome Analysis Reveal Recent Species-specific Gene Duplications in the Plastic Gilthead Sea Bream"

### Supplementary Figure 1

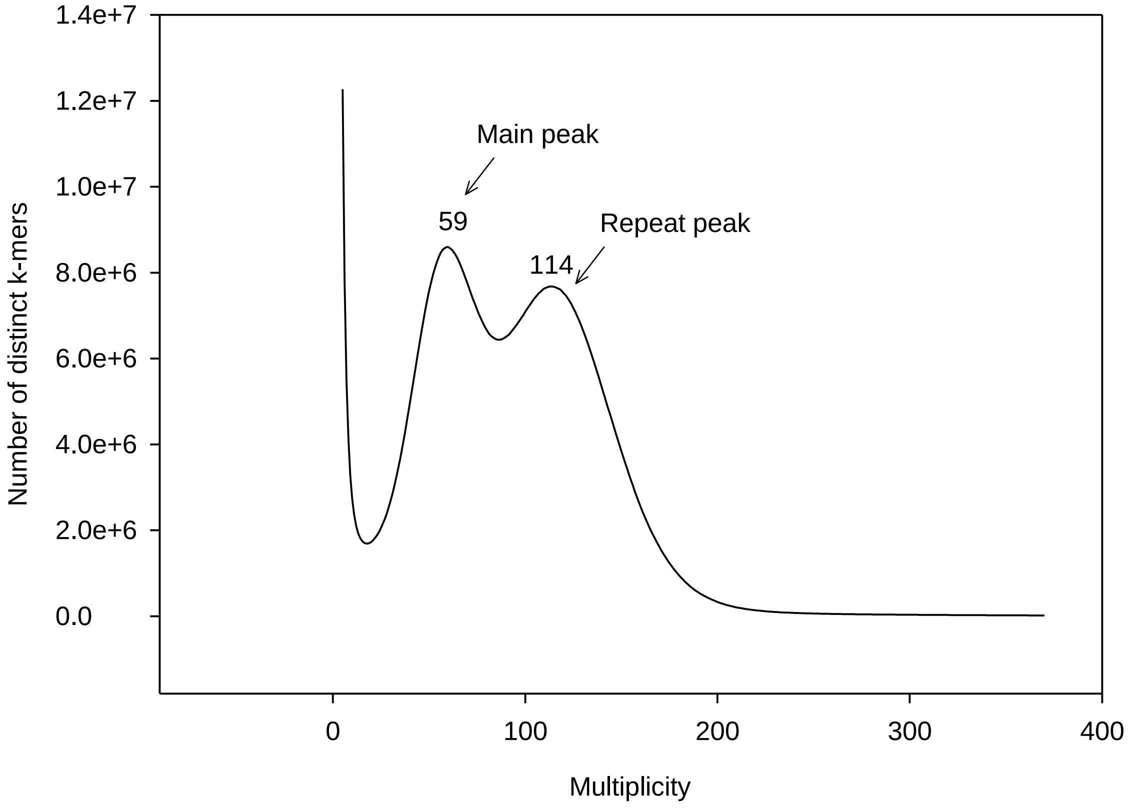

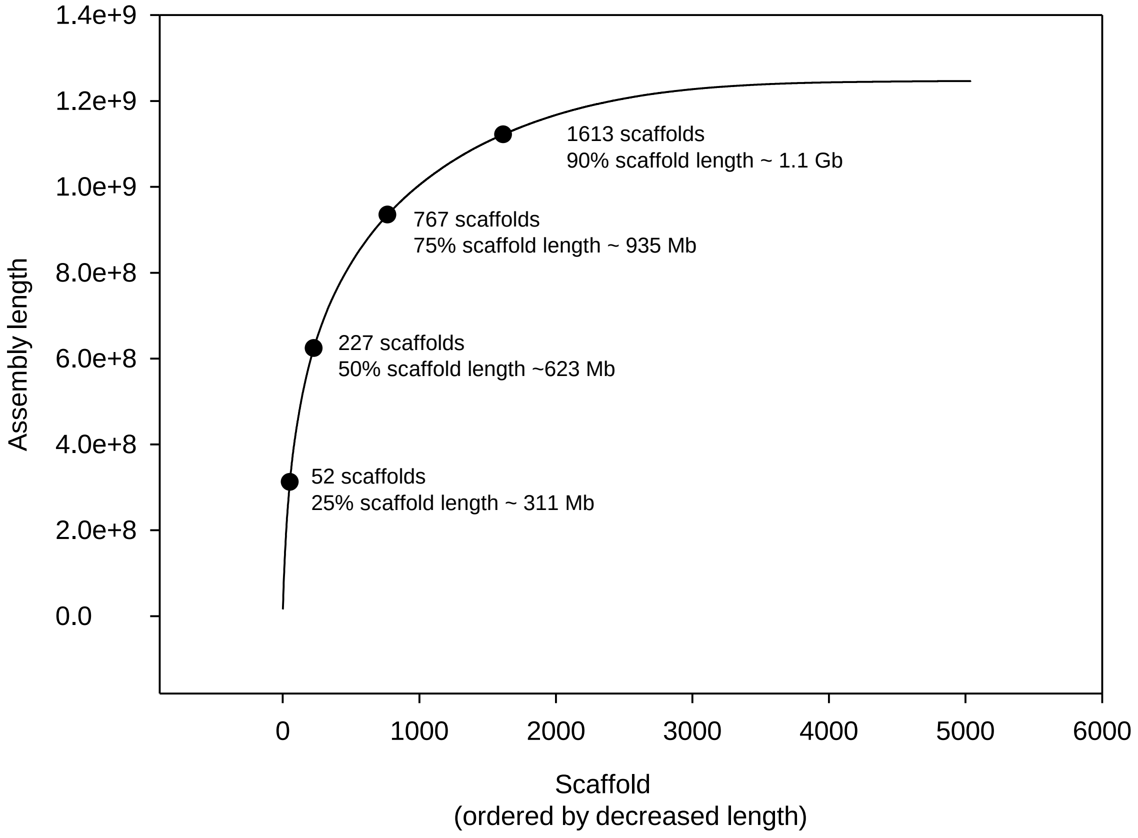

### Supplementary Figure 2

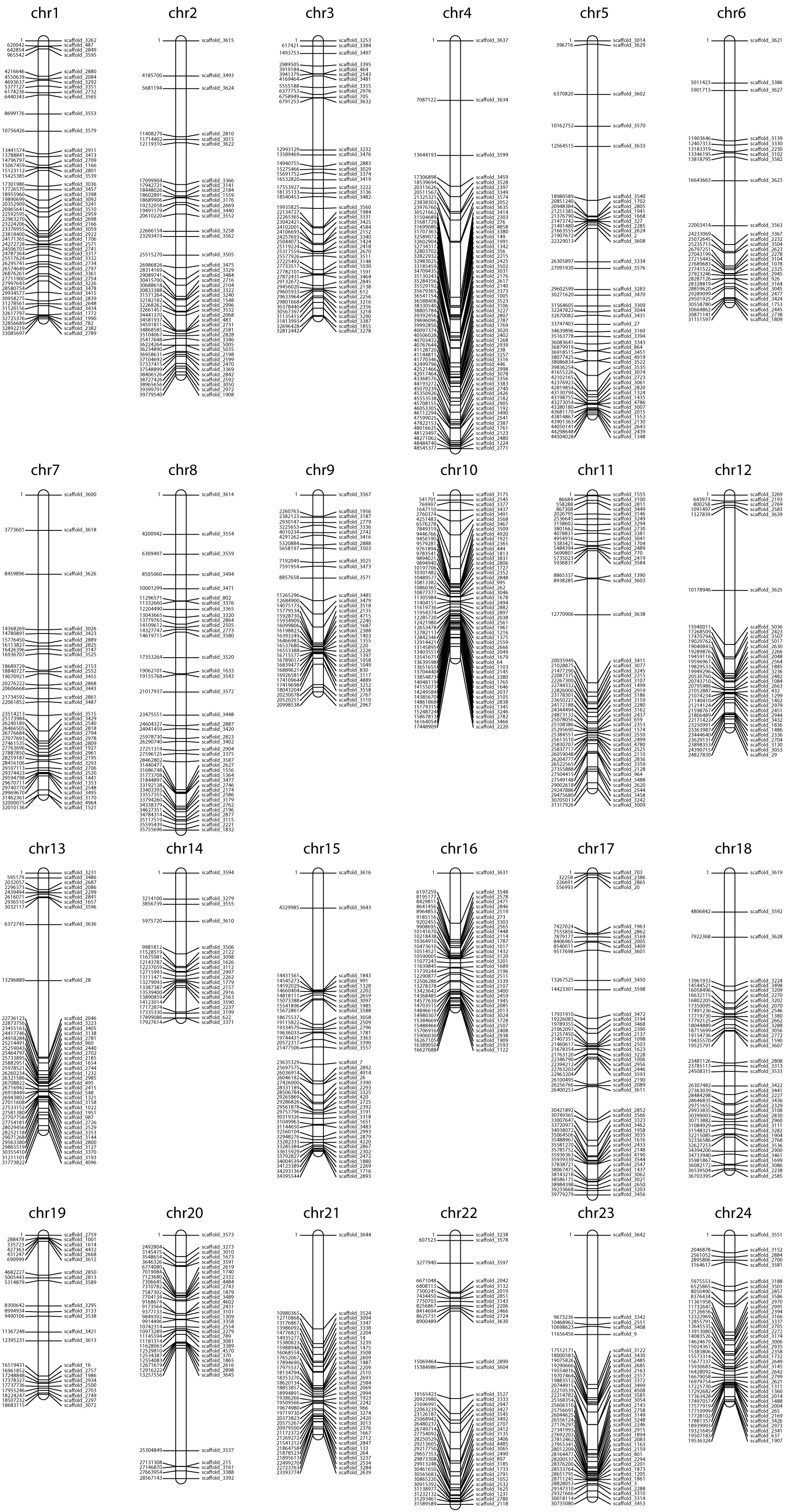
