## Supplementary Figure 3 for "Genome Sequencing and Transcriptome Analysis Reveal Recent Species-specific Gene Duplications in the Plastic Gilthead Sea Bream"

Javascript must be enabled to view this page.

magnitude
magnitudeUnassigned

04\_input\_krona\_classiffied\_krona

100.000001

53.840571

53.840570999

0.476487

0.007324

0.000954

0.001572

3.88523

0.044796

0.044424

0.021254

0.037365

0.011509

1.196957

0.098959

0.007006

0.349076

0.091804

0.000418

0.268835

0.000546

0.192949

10.050249

31.919118

0.001565

0.003028

0.149753

0.009976

0.014278

0.027208

4.927931

9.966608

9.966607999

0.097485

0.01265

0.02946

0.0019

0.000742

0.056244

0.015875

0.030014

0.008266

0.012744

0.011086

3.972008

0.029065

0.001514

0.000881

1.090599

0.000484

2.21887

0.019242

0.001465

0.025551

0.00041

0.022988

1.910308

0.01861

0.015131

0.220961

0.003491

0.01055

0.128014

6.369361

0.560519

0.016893

0.00023

0.031369

0.000173

0.003475

0.006461

0.0027

0.401786

0.002691

6.3e-05

0.094678

5.808842

0.000121

0.076902

0.071514

0.002299

0.30526

0.115032

0.003792

0.441944

4.77444

0.017538

29.823461

14.99056

0.247662

0.078201

0.001267

0.012993

0.000233

0.000124

0.000323

0.037403

12.425212

0.000683

0.006437

0.002966

0.006961

0.041155

2.12894

0.120121

0.120120999

14.71278

8.538525

0.190343

4.585233

0.645391

0.740747

0.012541
